# Identification of Catechins Binding Pockets in Monomeric A*β*_42_ Through Ensemble Docking and MD Simulations

**DOI:** 10.1101/2022.02.09.479729

**Authors:** Rohoullah Firouzi, Shahin Sowlati-Hashjin, Cecilia Chávez-García, Mitra Ashouri, Mohammad Hossein Karimi-Jafari, Mikko Karttunen

## Abstract

The assembly of the Amyloid-*β* peptide (A*β*) into toxic oligomers and fibrils is associated with Alzheimer’s disease and dementia. Therefore, disrupting amyloid assembly by direct targeting of the A*β* monomeric form with small molecules or antibodies is a promising therapeutic strategy. However, given the dynamic nature of A*β*, standard computational tools cannot be easily applied for high-throughput structure-based virtual screening in drug discovery projects. In the current study, we propose a computational pipeline – in the framework of the ensemble docking strategy – to identify catechins’ binding pockets in monomeric A*β*_42_. It is shown that both hydrophobic aromatic interactions and hydrogen bonding are crucial for the binding of catechins to A*β*_42_. Also, it has been found that all the studied ligands, especially the *EGCG*, can act as potent inhibitors against amyloid aggregation by blocking the central hydrophobic region of the A*β*. Our findings are evaluated and confirmed with multi-microsecond MD simulations. Finally, it is suggested that our proposed pipeline, with low computational cost in comparison with MD simulations, is a suitable approach for the virtual screening of ligand libraries against A*β*.

## 1. Introduction

Intrinsically disordered proteins (IDPs) are very flexible biomolecules without a well-defined folded structure and typically have important roles in biological processes, in particular in cellular signaling and gene regulation.^1–4^ Under certain conditions, some IDPs may aggregate into highly toxic oligomers. These oligomers are associated with a wide range of serious human diseases such as cancer, neurodegenerative diseases, autoimmune disorders, cardiovascular disease, and type II diabetes.^4–9^ Thus, preventing or reducing aggregation of the IDPs involved in such diseases appears as an effective therapeutic strategy.

In recent years, there have been efforts to design and synthesize small molecules and short peptides to block IDPs aggregation at different stages along the aggregation pathway, in particular nucleation and oligomer formation.^4,10–17^ Several studies have revealed that a large number of natural compounds derived from plants, animals and microorganisms, have the potential to inhibit oligomerization.^11–12, 18–20^ For example, several computational and experimental observations have shown that polyphenolic plant compounds which occur naturally in fruit, vegetables, chocolate, and tea are capable of inhibiting IDP aggregation.^12–13, 20–34^

Finding aggregation inhibitors and direct targeting of monomeric IDPs via small molecules is a very active area of research and a wide variety of computational techniques have been applied, yet there are many inherent difficulties.^12–14, 27, 35–41^ For example, most docking algorithms employ the flexible ligand and rigid receptor paradigm^42–44^ but IDPs display high conformational heterogeneity, and ligand binding causes large structural changes in the IDP conformations. To circumvent this problem, heterogeneous conformational ensembles of IDPs have been used for docking studies.^27, 38, 45–46^ Nevertheless, challenges associated with generating and choosing a set of suitable conformations for docking still remain. It will also be interesting to see how and if Alphafold^47^ will change the situation as it has already been applied to non-IDP related binding problems,^48^ but IDPs appear to be a challenge even for Alphafold.^49^

Several methods have been proposed for efficient sampling of IDP’s conformational space and constructing a representative conformational ensemble. Examples include replica-exchange-^50–53^ and metadynamics-based methods,^54–56^ diffusion map approaches^57–58^ and Markov state modeling.^59–62^ For comparisons between the methods, see for example Refs. ^39^ and ^63–65^. One alternative for effective exploration of the conformational space is the use of multiple conventional MD trajectories (replicas) with different initial conditions (different velocities or/and different starting configurations). The strategy of choosing the initial conditions controls the effectiveness of this approach and its ability to enhance conformational sampling performance.^66–70^ We have recently proposed a new efficient algorithm for comprehensive exploring of the conformational space of IDPs, called Blockwise Excursion Sampling (BES).^39^ It uses simulated annealing (SA) to find different low energy states of various regions of conformational space as optimal starting configurations for short conventional MD simulations. In BES, conformational sampling is based on many uncorrelated short MD simulations starting from different points of the accessible phase space. It has been shown that the protocol is successful in generating a diverse conformational library for IDP conformations in agreement with experimental data.^70^

In this work, we applied the BES protocol to generate a reliable structural library for the full-length human Amyloid-*β* (A*β*_42_) monomer involved in Alzheimer’s disease.^2,5,11–12,27,35,37,39^ Through ensemble docking approach, catechins, that are important natural polyphenolic compounds, were docked onto the surfaces to identify the binding “hot spots” on the A*β*_42_ monomers. It is noteworthy to point out that possible effects of catechins on A*β* aggregation have been the subject of numerous experimental and computational investigations. Those studies have proposed that catechins are able to disturb the A*β* aggregation and induce its aggregation into nontoxic forms oligomers.^21, 28,71–80^ In order to further evaluate the docking results and the stability of complexes, the obtained structures with the largest binding energies for each A*β*_42_-catechin complex were used as the starting structures for long multi-microsecond MD simulations (total of 15 μs).

## 2. Methods

### 2.1. Protein and Ligand Library Preparation

A library of A*β*_42_ (see **Figure 1** for a snapshot and the amino acid sequence) structures was generated using the BES protocol. Briefly, 2,000 excursion chains were performed such that all excursion chains were started from a fully extended structure; excursion chain refers to a sequence of MD and SA blocks in the BES protocol, for details and terminology please see Ref. ^70^.

**Figure 1.**
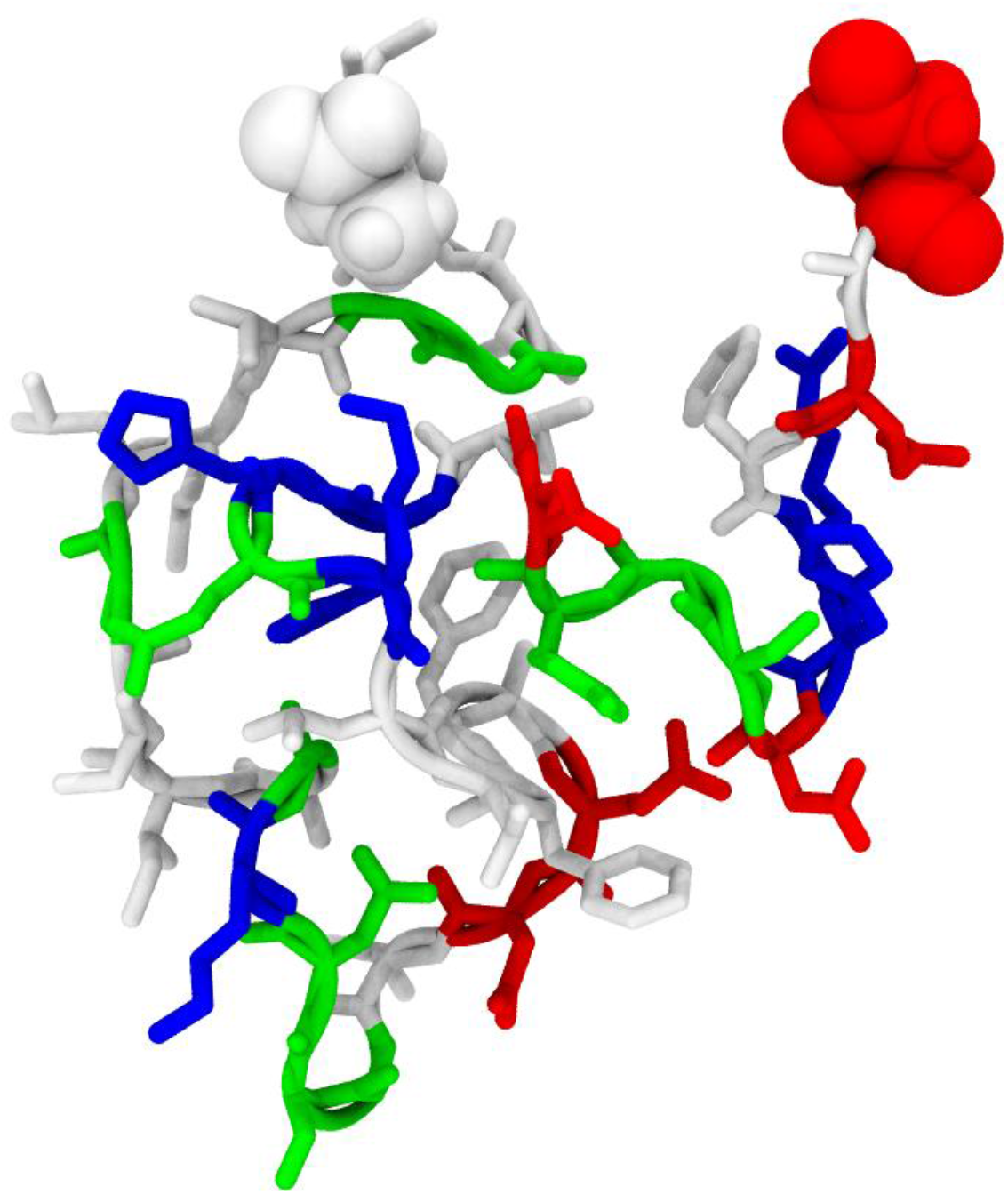
A random structure of Amyloid *β_42_*. A*β_42_* sequence: ^1^DAEFRHDSG ^10^YEVHHQKLVF ^20^FAEDVGSNKG ^30^AIIGLMVGGV ^40^VI^42^A. Asp1 (N-terminus) and Ala42 (C-terminus) are shown using van der Waals radii. Other residues are represented in licorice. Acidic, basic, polar, and non-polar amino acids are shown in red, blue, green, and white, respectively.

Each excursion chain included five successive SA and MD blocks with maximum temperatures of 700, 600, 500, 400, and 350 K for the SA blocks. The relaxation time for each MD block was set to 120 ps and the last 100 ps were used to generate representative structures. In the next step, an average (mean) structure over the MD trajectory was obtained, and the root mean deviation (RMSD) was used as a criterion to identify the configuration in the MD trajectory that is structurally closest to the average structure. The selected structure was then energy minimized using the conjugate gradient method and used as a representative structure. As a result, for each MD block (five blocks in each excursion chain) one representative structure was derived. The final structural library included a total of 10,000 representative structures. **Scheme 1** summarizes the procedure. For more technical details and a complete description of the BES protocol see Ref. ^70^.

**Scheme 1.**
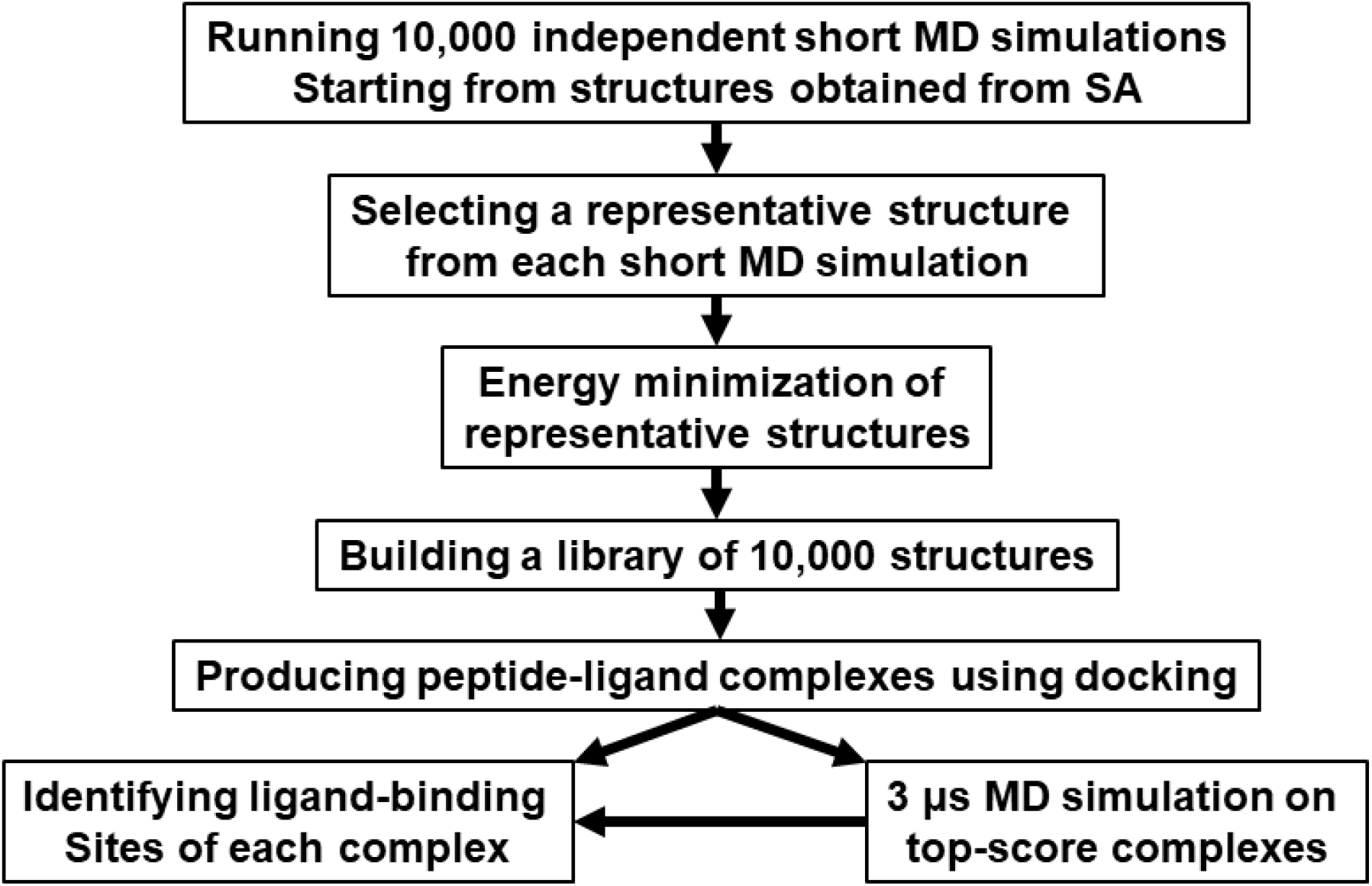
Flowchart of steps taken in this study.

Catechins (or Flavan-3-ols) are dietary polyphenolic compounds that many experimental results have suggested as potentially useful in targeting A*β*.^21, 28, 71–87^ For this study, we selected five different types of catechins: 1) (+)-catechin *(C),* 2) (-)-epicatechin (*EC*), 3) (-)-epigallocatechin (*EGC*), 4) (-)-epicatechin-3-gallate (*ECG*), and 5) (-)-epigallocatechin-3-gallate (*EGCG*). Their chemical structures are shown in **Figure 2** and their initial structures were taken from the ZINC database.^88^ This was followed by optimization of the molecular geometries for all of them using the B3LYP exchange and correlation functional^89^ and the 6-31+G(d,p) basis set. The GAMESS package^90^ was used for geometry optimization.

**Figure 2.**
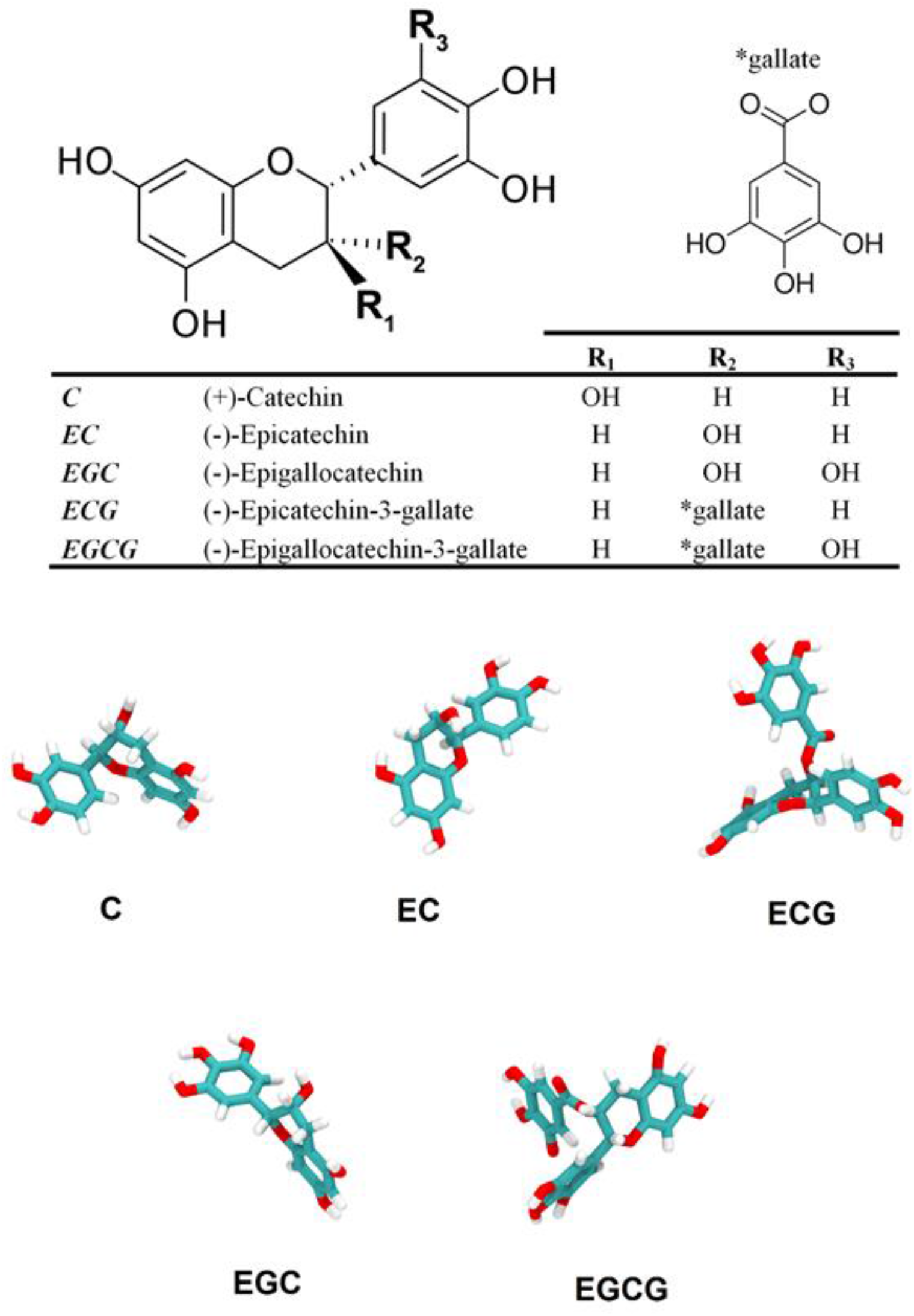
2D and 3D representations of catechins considered in this study.

### 2.2. Docking Setup

Docking simulations were performed with AutoDock Vina (version 1.1.2) software.^91^ The docking search space for exploring ligand binding conformations around each representative A*β*_42_ structure was defined using a rectangular box centered at the center of mass of A*β*_42_ with a minimal distance of 12 Å from the A*β*_42_ to the edges of the box. Therefore, depending on size and shape of the A*β*_42_ configuration, an optimized docking box was determined individually for each A*β*_42_. Each docking run generates nine optimal A*β*_42_-ligand bound conformations and overall, a total of 90,000 (10,000 × 9) poses were generated for each catechin compound. The different poses in each run are rank-ordered by the Vina score, a quantity that correlates with the binding free energy. The top-scoring pose in each run achieves the lowest free energy of binding in the complex.

Very recently, it has been shown that the correct pose (the pose with the lowest RMSD from the corresponding experimental pose) is usually predicted by Vina but sometimes, does not get the top score in the Vina ranking.^92–93^ To avoid the problem and to capture the correct poses, it is recommended that except for the top-ranked pose, some important lower ranked poses for each docking run should be identified and selected for post-docking analysis. For more discussions about ranking, see also Refs. ^94–97^. For this purpose, the differences in the binding free energies between the top-ranked pose and lower ranked poses were calculated for selected high-ranked modelled complexes for each docking run, *ΔΔG_binding_* (= ΔG_top-pose_ - ΔG_lower ranked pose_). Different cutoff values for the *ΔΔG_binding_* threshold (0.1, 0.2 and 0.3 kcal/mol) were used for selecting docking complexes, since the optimal selection of complexes is not known. For larger *ΔΔG_binding_* cutoff values, more complexes were selected. For example, with the cutoff value of 0.1 kcal/mol, 17,431 (10,000 × 1 (top-ranked poses from each run) + 7,431 (lower ranked poses near top-poses with the cutoff)) complexes were selected for docking of *EGCG*, while 29,923 and 42,833 complexes were selected with the cutoff values of 0.2 and 0.3 kcal/mol, respectively. The number of selected docking complexes for all ligands and the corresponding cutoff values of *ΔΔG_binding_* are provided and labelled as “set-*n*” in **Table 1**. Since the results for different sets are very similar, we only present the results of set-1 (with the cutoff value of 0.1) and the results for the other two sets can be found in Supporting Information.

**Table 1.**
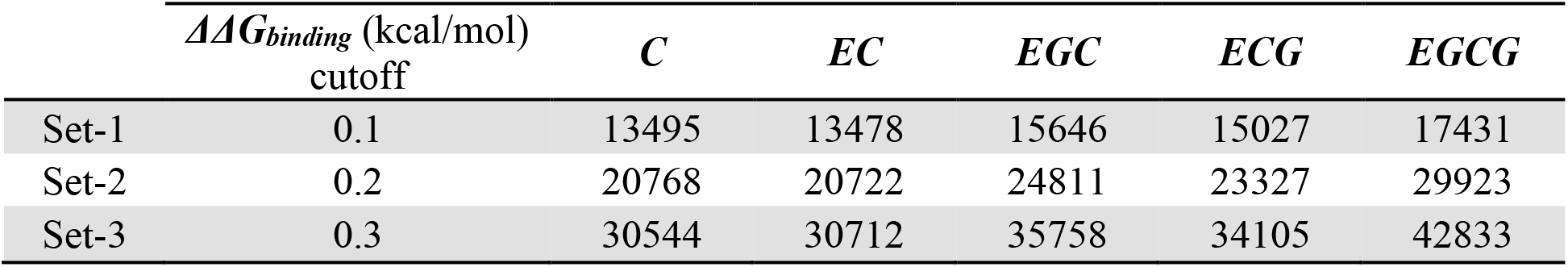
The number of selected docking complexes for each ligand with the different *ΔΔG_binding_* cutoffs.

### 2.3. MD simulation setup

All-atom MD simulations were performed on the peptide with five different ligands. The force field parameters for each ligand were created using the Antechamber program in the Ambertools19 package^98^ and described by the General Amber Force Field (GAFF)^99^ using AM1-BCC charges.^100^ The Amberff99SB*-ILDNP force field^101^ and the TIP3P water model^102^ were adopted for the protein and water, respectively. Each protein was placed in a dodecahedral box such that the distance from the edges of the box to every atom in the protein was at least 1 nm and 150 mM of KCl was added to reproduce physiological conditions. Overall charge neutrality was preserved by adding 3 K^+^ counterions. The GROMACS 2016.3^103^ package was used for all simulations. Each system was energy minimized using the method of steepest descents. This was followed by a pre-equilibration in the canonical ensemble, i.e., at constant particle number, volume, and temperature, for 100 ps. The Lennard-Jones potential was truncated using a shift function between 1.0 and 1.2 nm. Electrostatic interactions were calculated using the particle-mesh Ewald method (PME)^104–105^ with a real space cut-off of 1.2 nm. The temperature was set to 310 K with the V-rescale algorithm^106^ and pressure was kept at 1 atm using the Parrinello-Rahman barostat.^107^ Bonds involving hydrogens were constrained using the Linear Constraint Solver (P-LINCS) algorithm.^108^ Pre-equilibration was followed by a production run of 3 μs with a time step of 2 fs for each of the five peptide-ligand systems.

## 3. Results and Discussion

### 3.1. Docking Analysis

#### Structural Analysis and Identifying Ligand Binding Site

Determination of the important residues that are in close contact with the ligand is very important for the identification of potential inhibitors of A*β*_42_ aggregation. The distances between the heavy atoms of the ligand and the residues of A*β*_42_ were used to define the binding sites of A*β*_42_. Based on this assumption, a list of binding residues for each selected complex was generated for which at least one heavy atom of the residues falls within the cutoff distance of any ligand heavy atom. To assess the effect of the cutoff distance, five distances were evaluated: 3.0, 3.5, 4.0, 4.5, and 5.0 Å; cutoff distances from 3.0 to 5.0 Å are commonly used to study ligand-protein binding interactions, such as hydrogen bonds, hydrophobic contacts, and aromatic interactions.^22, 109–114^

The number of contacts between each ligand and A*β*_42_ residues for the set-1 (with *ΔΔG_binding_* = 0.1 kcal/mol) was counted, see **Tables 2-6**. The first observation from the tables is that all the ligands have the most contacts with residues Tyr10, Phe19, and Phe20. Moreover, based on the ranking (**Tables 1-6**), other aromatic residues of (His13, His14, His6, and Phe4) also contribute to stabilizing the interactions with catechins: These polyphenolic compounds tend to interact with the aromatic residues through stacking and/or T-shaped interactions, see the snapshots in **Figures 3-7**. Another important observation is that the tendency of ligands to interact with Tyr10 correlates with the number of hydroxyl groups on the ligands. This is seen in the data in **Tables 2-6**: *EGCG* possesses the largest number of hydroxyl groups (8 OH’s) and has a greater tendency to interact with Tyr10 than with Phe19 or Phe20, i.e., the number of contacts between *EGCG* and Tyr10 is larger than those between *EGCG* and Phe19 or Phe20. Thus, the data appears to imply that, hydrophobic aromatic interactions and hydrogen bonding are both crucial for the binding process. Finally, a comparative look at the tabulated values immediately shows that all ligands have a tendency to associate with the hydrophobic region of A*β*_42_ spanning residues from Tyr10 to Phe20. This region contains most of the aromatic residues found in full-length A*β*. This region also encompasses the central hydrophobic region (^16^KLVFF^20^) that, based on many experimental and computational studies, is involved in the initiation of amyloid aggregation.^11–13, 22, 36, 115–119^ Therefore, our docking results show that all studied ligands, especially the *EGCG*, can act as potent inhibitors against amyloid aggregation through blocking the central hydrophobic region. These findings are in agreement with experimental studies.^28, 72–74^

**Table 2.** The number of contacts between the amino acid residues (the amino acid sequence is provided in the caption of **Figure 1**) of A*β*_42_ and the *C* ligand of set-1 (13,495 complexes, **Table 1**) with five different distance cutoffs as indicated. The aromatic residues are in bold typeface and three most favorable aromatic residue hotspots are highlighted in light teal.

| 3.0 Å |  |  | 3.5 Å |  |  | 4.0 Å |  |  | 4.5 Å |  |  | 5.0 Å |  |  |
| --- | --- | --- | --- | --- | --- | --- | --- | --- | --- | --- | --- | --- | --- | --- |
| Residue | ID | Population | Residue | ID | Population | Residue | ID | Population | Residue | ID | Population | Residue | ID | Population |
| **PHE** | **19** | **4455** | **PHE** | **19** | **5160** | **PHE** | **19** | **5510** | **PHE** | **19** | **5853** | **PHE** | **19** | **6290** |
| **TYR** | **10** | **4263** | **TYR** | **10** | **4916** | **PHE** | **20** | **5297** | **PHE** | **20** | **5739** | **PHE** | **20** | **6242** |
| **PHE** | **20** | **4138** | **PHE** | **20** | **4884** | **TYR** | **10** | **5267** | **TYR** | **10** | **5603** | **TYR** | **10** | **6014** |
| GLN | 15 | 3963 | GLN | 15 | 4602 | GLN | 15 | 4987 | GLN | 15 | 5419 | GLN | 15 | 5911 |
| LYS | 16 | 3745 | LYS | 16 | 4220 | LYS | 16 | 4611 | LYS | 16 | 5040 | LYS | 16 | 5586 |
| LEU | 17 | 3472 | LEU | 17 | 4053 | LEU | 17 | 4515 | LEU | 17 | 4999 | LEU | 17 | 5581 |
| **HIS** | **13** | **3402** | **HIS** | **13** | **4030** | **HIS** | **13** | **4441** | **HIS** | **13** | **4885** | VAL | 18 | 5423 |
| **HIS** | **14** | **3338** | **HIS** | **14** | **4000** | **HIS** | **14** | **4412** | **HIS** | **14** | **4850** | **HIS** | **13** | **5386** |
| VAL | 18 | 3246 | VAL | 18 | 3938 | VAL | 18 | 4371 | VAL | 18 | 4838 | **HIS** | **14** | **5381** |
| VAL | 12 | 3027 | VAL | 12 | 3713 | VAL | 12 | 4171 | VAL | 12 | 4661 | VAL | 12 | 5279 |
| GLU | 11 | 2926 | GLU | 11 | 3418 | GLU | 11 | 3797 | GLU | 11 | 4265 | GLU | 11 | 4785 |
| ARG | 5 | 2642 | ALA | 21 | 3237 | ALA | 21 | 3688 | ALA | 21 | 4169 | ALA | 21 | 4721 |
| **PHE** | **4** | **2628** | GLU | 22 | 3061 | GLU | 22 | 3411 | GLU | 22 | 3776 | GLY | 9 | 4225 |
| ALA | 21 | 2617 | ARG | 5 | 3054 | ARG | 5 | 3360 | GLY | 9 | 3718 | GLU | 22 | 4223 |
| GLU | 22 | 2587 | **HIS** | **6** | **3038** | **HIS** | **6** | **3321** | ARG | 5 | 3712 | SER | 8 | 4122 |
| **HIS** | **6** | **2530** | **PHE** | **4** | **3026** | GLY | 9 | 3262 | SER | 8 | 3632 | ARG | 5 | 4075 |
| SER | 8 | 2337 | SER | 8 | 2869 | SER | 8 | 3258 | **HIS** | **6** | **3615** | **HIS** | **6** | **3934** |
| GLY | 9 | 2265 | GLY | 9 | 2866 | **PHE** | **4** | **3255** | **PHE** | **4** | **3452** | ASP | 7 | 3715 |
| VAL | 24 | 2154 | ASP | 7 | 2587 | VAL | 24 | 2931 | VAL | 24 | 3289 | VAL | 24 | 3710 |
| ASP | 7 | 2140 | VAL | 24 | 2587 | ASP | 7 | 2904 | ASP | 7 | 3265 | **PHE** | **4** | **3686** |
| ASP | 23 | 1905 | ASP | 23 | 2327 | ASP | 23 | 2602 | ASP | 23 | 2985 | ASP | 23 | 3430 |
| ASN | 27 | 1895 | ASN | 27 | 2190 | ASN | 27 | 2393 | SER | 26 | 2721 | SER | 26 | 3110 |
| SER | 26 | 1738 | SER | 26 | 2076 | SER | 26 | 2374 | ASN | 27 | 2667 | ASN | 27 | 2975 |
| GLU | 3 | 1705 | GLU | 3 | 2037 | GLU | 3 | 2256 | GLY | 25 | 2514 | GLY | 25 | 2927 |
| LYS | 28 | 1684 | LYS | 28 | 1957 | GLY | 25 | 2186 | GLU | 3 | 2463 | LYS | 28 | 2766 |
| GLY | 25 | 1459 | GLY | 25 | 1879 | LYS | 28 | 2172 | LYS | 28 | 2433 | GLU | 3 | 2685 |
| ILE | 31 | 1452 | ILE | 31 | 1711 | ILE | 31 | 1916 | ILE | 31 | 2172 | ILE | 31 | 2473 |
| ALA | 30 | 1327 | ALA | 30 | 1613 | ALA | 30 | 1844 | ALA | 30 | 2101 | ALA | 30 | 2427 |
| ILE | 32 | 1291 | ILE | 32 | 1538 | ILE | 32 | 1749 | GLY | 29 | 1981 | GLY | 29 | 2310 |
| LEU | 34 | 1225 | GLY | 29 | 1489 | GLY | 29 | 1719 | ILE | 32 | 1979 | ILE | 32 | 2254 |
| ALA | 2 | 1169 | LEU | 34 | 1461 | ALA | 2 | 1641 | ALA | 2 | 1843 | LEU | 34 | 2110 |
| GLY | 29 | 1151 | ALA | 2 | 1434 | LEU | 34 | 1637 | LEU | 34 | 1842 | ALA | 2 | 2040 |
| VAL | 36 | 1049 | VAL | 36 | 1264 | MET | 35 | 1449 | MET | 35 | 1652 | MET | 35 | 1875 |
| MET | 35 | 966 | MET | 35 | 1245 | VAL | 36 | 1415 | VAL | 36 | 1582 | VAL | 36 | 1822 |
| GLY | 33 | 914 | GLY | 33 | 1148 | GLY | 33 | 1297 | GLY | 33 | 1471 | GLY | 33 | 1672 |
| VAL | 39 | 854 | VAL | 39 | 1048 | VAL | 39 | 1193 | VAL | 39 | 1342 | VAL | 39 | 1559 |
| VAL | 40 | 832 | VAL | 40 | 982 | ASP | 1 | 1116 | ASP | 1 | 1265 | ASP | 1 | 1416 |
| ASP | 1 | 806 | ASP | 1 | 977 | VAL | 40 | 1084 | VAL | 40 | 1225 | VAL | 40 | 1392 |
| ILE | 41 | 771 | ILE | 41 | 948 | ILE | 41 | 1074 | ILE | 41 | 1213 | GLY | 37 | 1373 |
| GLY | 37 | 648 | GLY | 37 | 815 | GLY | 37 | 984 | GLY | 37 | 1135 | ILE | 41 | 1372 |
| GLY | 38 | 643 | GLY | 38 | 814 | GLY | 38 | 953 | GLY | 38 | 1092 | GLY | 38 | 1305 |
| ALA | 42 | 468 | ALA | 42 | 580 | ALA | 42 | 674 | ALA | 42 | 764 | ALA | 42 | 870 |

**Table 3.**
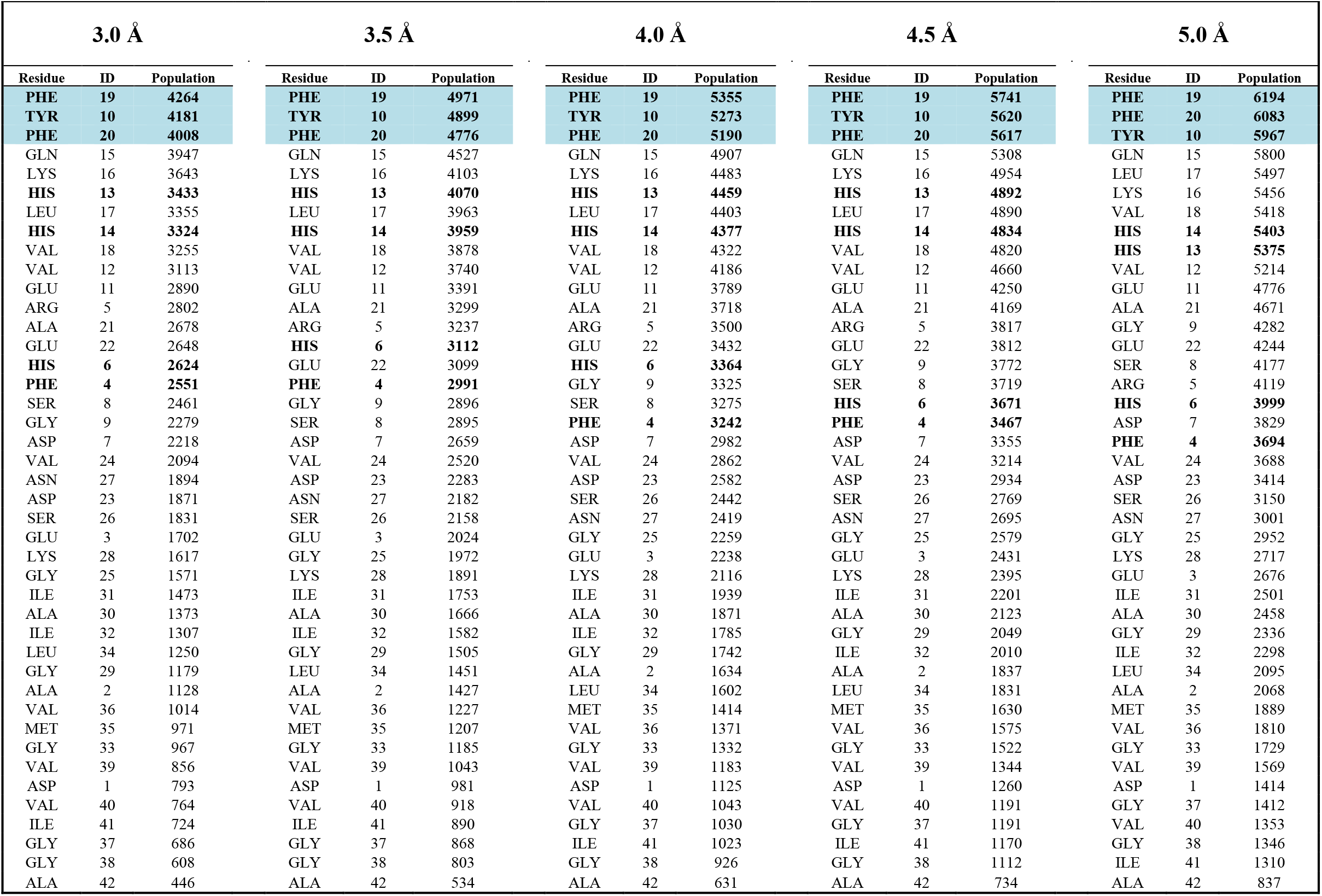
The number of contacts between the amino acid residues (the amino acid sequence is provided in the caption of **Figure 1**) of A*β*_42_ and the *EC* ligand for set-1 (13,478 complexes, **Table 1**) with five different distance cutoffs as indicated. The aromatic residues are in bold typeface and three most favorable aromatic residue hotspots are highlighted in light teal.

**Table 4.** The number of contacts between the amino acid residues (the amino acid sequence is provided in the caption of **Figure 1**) of A*β*_42_ and the *ECG* ligand for set-1 (15,027 complexes, **Table 1**) with five different distance cutoffs as indicated. The aromatic residues are in bold typeface and three most favorable aromatic residue hotspots are highlighted in light teal.

| 3.0 Å |  |  | 3.5 Å |  |  | 4.0 Å |  |  | 4.5 Å |  |  | 5.0 Å |  |  |
| --- | --- | --- | --- | --- | --- | --- | --- | --- | --- | --- | --- | --- | --- | --- |
| Residue | ID | Population | Residue | ID | Population | Residue | ID | Population | Residue | ID | Population | Residue | ID | Population |
| **PHE** | **19** | **5225** | **PHE** | **19** | **6050** | **PHE** | **19** | **6491** | **PHE** | **19** | **6953** | **PHE** | **19** | **7425** |
| **TYR** | **10** | **5192** | **TYR** | **10** | **5937** | **TYR** | **10** | **6338** | **PHE** | **20** | **6751** | **PHE** | **20** | **7304** |
| **PHE** | **20** | **4974** | **PHE** | **20** | **5792** | **PHE** | **20** | **6266** | **TYR** | **10** | **6743** | **TYR** | **10** | **7187** |
| GLN | 15 | 4954 | GLN | 15 | 5629 | GLN | 15 | 6025 | GLN | 15 | 6506 | GLN | 15 | 7058 |
| LYS | 16 | 4384 | **HIS** | **13** | **5062** | **HIS** | **13** | **5589** | **HIS** | **13** | **6053** | **HIS** | **14** | **6641** |
| **HIS** | **13** | **4322** | LYS | 16 | 4985 | LYS | 16 | 5467 | **HIS** | **14** | **6017** | **HIS** | **13** | **6620** |
| **HIS** | **14** | **4218** | **HIS** | **14** | **4944** | **HIS** | **14** | **5459** | LYS | 16 | 5968 | LEU | 17 | 6578 |
| LEU | 17 | 4134 | LEU | 17 | 4838 | LEU | 17 | 5332 | LEU | 17 | 5901 | VAL | 18 | 6545 |
| VAL | 18 | 4081 | VAL | 18 | 4820 | VAL | 18 | 5322 | VAL | 18 | 5877 | LYS | 16 | 6545 |
| VAL | 12 | 4029 | VAL | 12 | 4770 | VAL | 12 | 5305 | VAL | 12 | 5857 | VAL | 12 | 6495 |
| GLU | 11 | 3786 | GLU | 11 | 4406 | GLU | 11 | 4862 | GLU | 11 | 5393 | GLU | 11 | 5988 |
| ARG | 5 | 3652 | ALA | 21 | 4112 | ALA | 21 | 4620 | ALA | 21 | 5151 | ALA | 21 | 5763 |
| ALA | 21 | 3416 | ARG | 5 | 4101 | ARG | 5 | 4460 | GLY | 9 | 4876 | GLY | 9 | 5452 |
| GLU | 22 | 3332 | GLU | 22 | 3894 | GLY | 9 | 4328 | ARG | 5 | 4796 | SER | 8 | 5246 |
| **HIS** | **6** | **3297** | SER | 8 | 3889 | SER | 8 | 4312 | SER | 8 | 4712 | ARG | 5 | 5178 |
| SER | 8 | 3275 | **HIS** | **6** | **3882** | GLU | 22 | 4260 | GLU | 22 | 4690 | GLU | 22 | 5176 |
| **PHE** | **4** | **3272** | GLY | 9 | 3830 | **HIS** | **6** | **4245** | **HIS** | **6** | **4610** | **HIS** | **6** | **4978** |
| GLY | 9 | 3080 | **PHE** | **4** | **3787** | **PHE** | **4** | **4088** | **PHE** | **4** | **4357** | ASP | 7 | 4792 |
| ASP | 7 | 2965 | ASP | 7 | 3563 | ASP | 7 | 3917 | ASP | 7 | 4315 | **PHE** | **4** | **4623** |
| VAL | 24 | 2712 | VAL | 24 | 3259 | VAL | 24 | 3632 | VAL | 24 | 4062 | VAL | 24 | 4612 |
| ASP | 23 | 2550 | ASP | 23 | 3042 | ASP | 23 | 3404 | ASP | 23 | 3805 | ASP | 23 | 4334 |
| ASN | 27 | 2459 | ASN | 27 | 2851 | SER | 26 | 3202 | SER | 26 | 3560 | SER | 26 | 3998 |
| SER | 26 | 2413 | SER | 26 | 2848 | ASN | 27 | 3116 | ASN | 27 | 3406 | ASN | 27 | 3808 |
| GLU | 3 | 2320 | GLU | 3 | 2752 | GLU | 3 | 3025 | GLY | 25 | 3273 | GLY | 25 | 3751 |
| LYS | 28 | 2143 | GLY | 25 | 2527 | GLY | 25 | 2890 | GLU | 3 | 3259 | GLU | 3 | 3548 |
| GLY | 25 | 2052 | LYS | 28 | 2468 | LYS | 28 | 2772 | LYS | 28 | 3153 | LYS | 28 | 3512 |
| ILE | 31 | 1780 | ILE | 31 | 2114 | ILE | 31 | 2365 | ILE | 31 | 2642 | ALA | 30 | 3044 |
| ALA | 30 | 1659 | ALA | 30 | 2066 | ALA | 30 | 2325 | ALA | 30 | 2634 | ILE | 31 | 3014 |
| ILE | 32 | 1640 | GLY | 29 | 1937 | GLY | 29 | 2248 | GLY | 29 | 2533 | GLY | 29 | 2903 |
| LEU | 34 | 1568 | ILE | 32 | 1932 | ALA | 2 | 2194 | ILE | 32 | 2476 | ILE | 32 | 2808 |
| ALA | 2 | 1529 | ALA | 2 | 1919 | ILE | 32 | 2191 | ALA | 2 | 2470 | ALA | 2 | 2753 |
| GLY | 29 | 1507 | LEU | 34 | 1841 | LEU | 34 | 2027 | LEU | 34 | 2252 | LEU | 34 | 2549 |
| VAL | 36 | 1310 | VAL | 36 | 1589 | MET | 35 | 1803 | MET | 35 | 2060 | MET | 35 | 2415 |
| MET | 35 | 1251 | MET | 35 | 1553 | VAL | 36 | 1786 | VAL | 36 | 2018 | VAL | 36 | 2284 |
| ASP | 1 | 1192 | GLY | 33 | 1436 | GLY | 33 | 1646 | GLY | 33 | 1854 | GLY | 33 | 2139 |
| GLY | 33 | 1163 | ASP | 1 | 1435 | ASP | 1 | 1618 | ASP | 1 | 1805 | VAL | 39 | 2038 |
| VAL | 39 | 1115 | VAL | 39 | 1347 | VAL | 39 | 1572 | VAL | 39 | 1785 | ASP | 1 | 2008 |
| ILE | 41 | 1046 | VAL | 40 | 1267 | VAL | 40 | 1424 | ILE | 41 | 1622 | VAL | 40 | 1824 |
| VAL | 40 | 1043 | ILE | 41 | 1234 | ILE | 41 | 1420 | VAL | 40 | 1604 | ILE | 41 | 1805 |
| GLY | 37 | 848 | GLY | 37 | 1094 | GLY | 37 | 1300 | GLY | 37 | 1505 | GLY | 37 | 1792 |
| GLY | 38 | 836 | GLY | 38 | 1073 | GLY | 38 | 1272 | GLY | 38 | 1477 | GLY | 38 | 1761 |
| ALA | 42 | 656 | ALA | 42 | 829 | ALA | 42 | 948 | ALA | 42 | 1082 | ALA | 42 | 1202 |

**Table 5.**
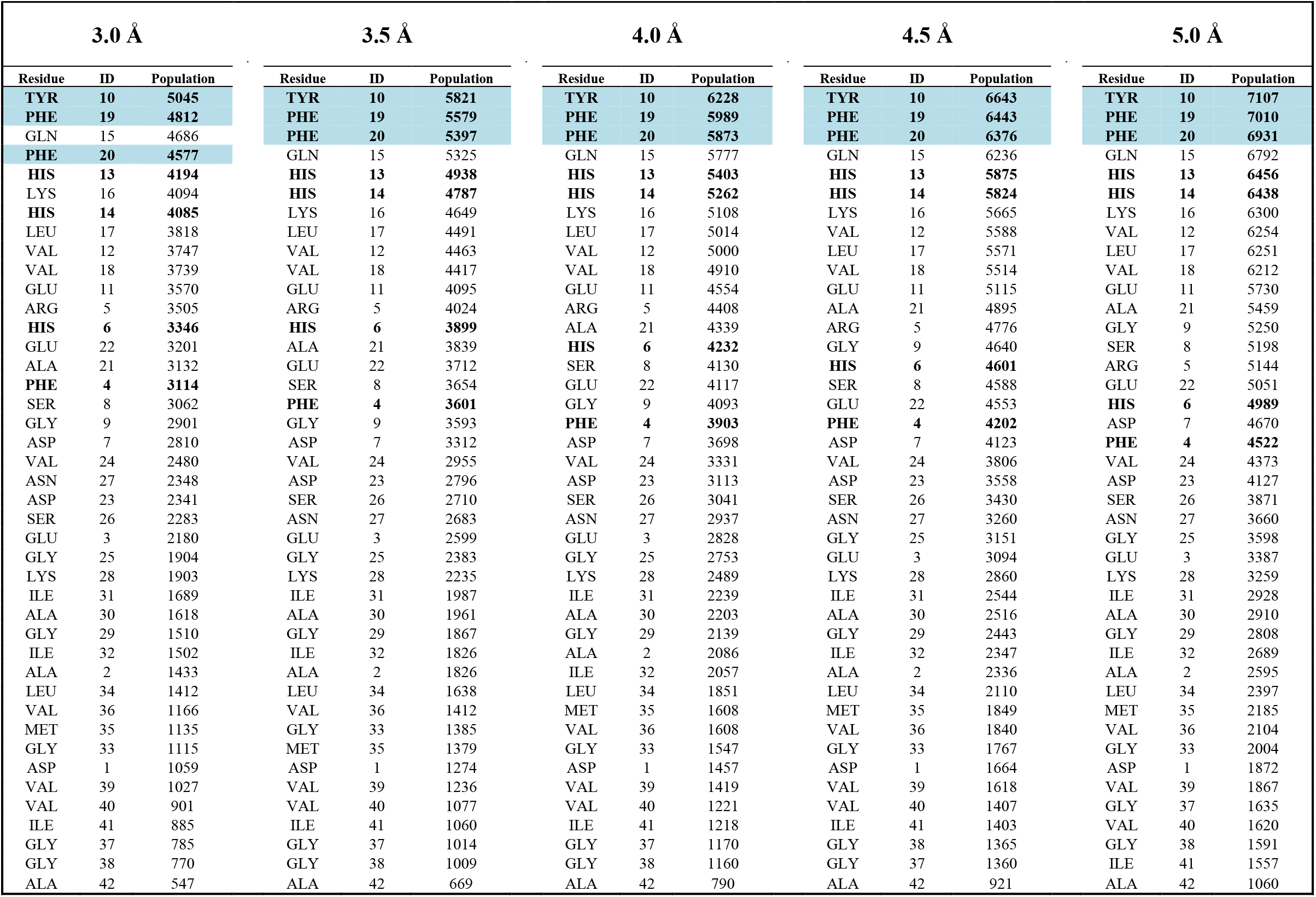
The number of contacts between the amino acid residues (the amino acid sequence is provided in the caption of **Figure 1**) of A*β*_42_ and the *EGC* ligand for set-1 (15,646 complexes, **Table 1**) with five different distance cutoffs as indicated. The aromatic residues are in bold typeface and the three most favorable aromatic residue hotspots are highlighted in light teal.

**Table 6.**
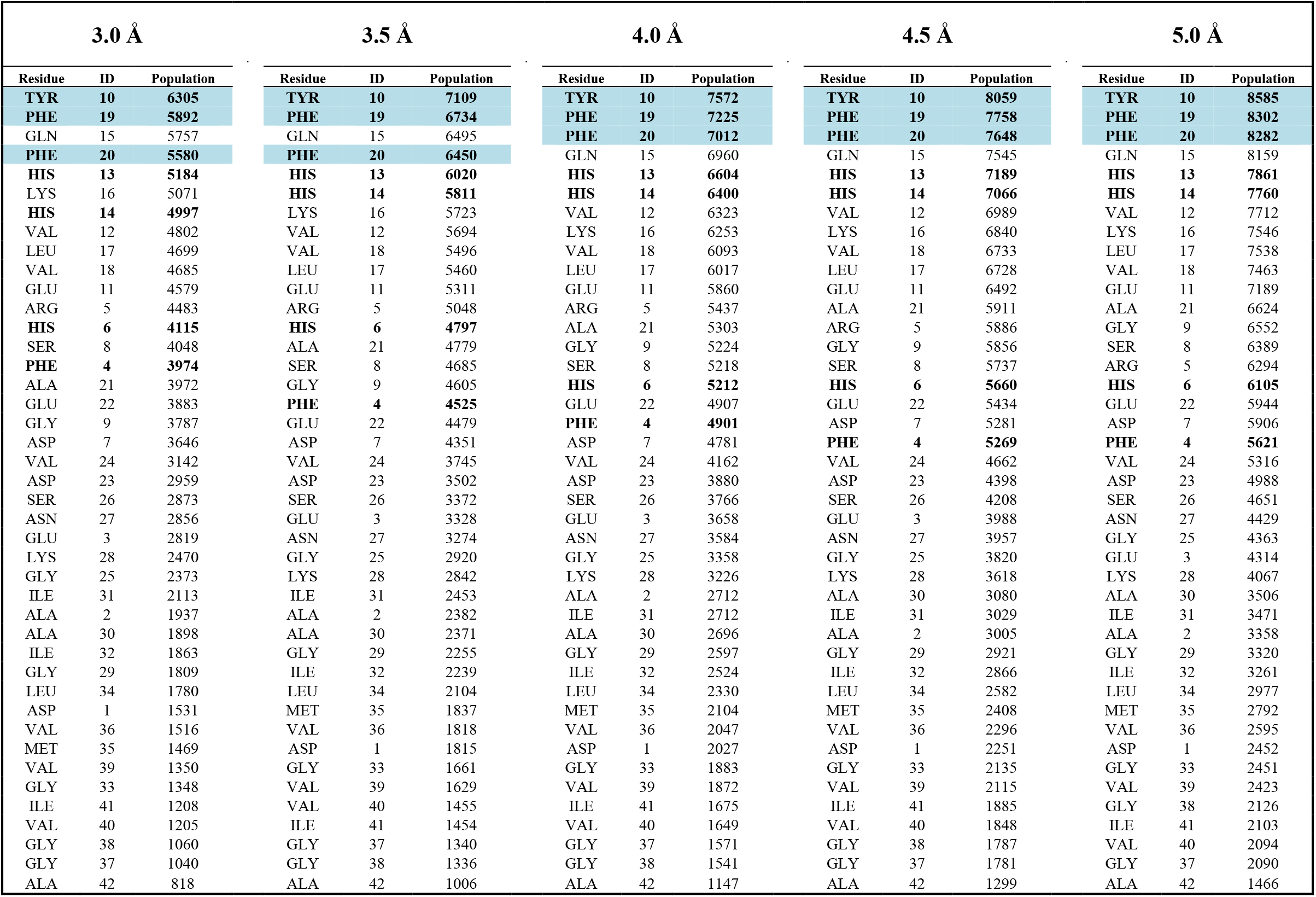
The number of contacts between the amino acid residues (the amino acid sequence is provided in the caption of **Figure 1**) of A*β*_42_ and the *EGCG* ligand for set-1 (17,431 complexes, **Table 1**) with five different distance cutoffs as indicated. The aromatic residues are in bold typeface and three most favorable aromatic residue hotspots are highlighted in light teal.

| 3.0 Å |  |  | 3.5 Å |  |  | 4.0 Å |  |  | 4.5 Å |  |  | 5.0 Å |  |  |
| --- | --- | --- | --- | --- | --- | --- | --- | --- | --- | --- | --- | --- | --- | --- |
| Residue | ID | Population | Residue | ID | Population | Residue | ID | Population | Residue | ID | Population | Residue | ID | Population |
| **TYR** | **10** | **6305** | **TYR** | **10** | **7109** | **TYR** | **10** | **7572** | **TYR** | **10** | **8059** | **TYR** | **10** | **8585** |
| **PHE** | **19** | **5892** | **PHE** | **19** | **6734** | **PHE** | **19** | **7225** | **PHE** | **19** | **7758** | **PHE** | **19** | **8302** |
| GLN | 15 | 5757 | GLN | 15 | 6495 | **PHE** | **20** | **7012** | **PHE** | **20** | **7648** | **PHE** | **20** | **8282** |
| **PHE** | **20** | **5580** | **PHE** | **20** | **6450** | GLN | 15 | 6960 | GLN | 15 | 7545 | GLN | 15 | 8159 |
| **HIS** | **13** | **5184** | **HIS** | **13** | **6020** | **HIS** | **13** | **6604** | **HIS** | **13** | **7189** | **HIS** | **13** | **7861** |
| LYS | 16 | 5071 | **HIS** | **14** | **5811** | **HIS** | **14** | **6400** | **HIS** | **14** | **7066** | **HIS** | **14** | **7760** |
| **HIS** | **14** | **4997** | LYS | 16 | 5723 | VAL | 12 | 6323 | VAL | 12 | 6989 | VAL | 12 | 7712 |
| VAL | 12 | 4802 | VAL | 12 | 5694 | LYS | 16 | 6253 | LYS | 16 | 6840 | LYS | 16 | 7546 |
| LEU | 17 | 4699 | VAL | 18 | 5496 | VAL | 18 | 6093 | VAL | 18 | 6733 | LEU | 17 | 7538 |
| VAL | 18 | 4685 | LEU | 17 | 5460 | LEU | 17 | 6017 | LEU | 17 | 6728 | VAL | 18 | 7463 |
| GLU | 11 | 4579 | GLU | 11 | 5311 | GLU | 11 | 5860 | GLU | 11 | 6492 | GLU | 11 | 7189 |
| ARG | 5 | 4483 | ARG | 5 | 5048 | ARG | 5 | 5437 | ALA | 21 | 5911 | ALA | 21 | 6624 |
| **HIS** | **6** | **4115** | **HIS** | **6** | **4797** | ALA | 21 | 5303 | ARG | 5 | 5886 | GLY | 9 | 6552 |
| SER | 8 | 4048 | ALA | 21 | 4779 | GLY | 9 | 5224 | GLY | 9 | 5856 | SER | 8 | 6389 |
| **PHE** | **4** | **3974** | SER | 8 | 4685 | SER | 8 | 5218 | SER | 8 | 5737 | ARG | 5 | 6294 |
| ALA | 21 | 3972 | GLY | 9 | 4605 | **HIS** | **6** | **5212** | **HIS** | **6** | **5660** | **HIS** | **6** | **6105** |
| GLU | 22 | 3883 | **PHE** | **4** | **4525** | GLU | 22 | 4907 | GLU | 22 | 5434 | GLU | 22 | 5944 |
| GLY | 9 | 3787 | GLU | 22 | 4479 | **PHE** | **4** | **4901** | ASP | 7 | 5281 | ASP | 7 | 5906 |
| ASP | 7 | 3646 | ASP | 7 | 4351 | ASP | 7 | 4781 | **PHE** | **4** | **5269** | **PHE** | **4** | **5621** |
| VAL | 24 | 3142 | VAL | 24 | 3745 | VAL | 24 | 4162 | VAL | 24 | 4662 | VAL | 24 | 5316 |
| ASP | 23 | 2959 | ASP | 23 | 3502 | ASP | 23 | 3880 | ASP | 23 | 4398 | ASP | 23 | 4988 |
| SER | 26 | 2873 | SER | 26 | 3372 | SER | 26 | 3766 | SER | 26 | 4208 | SER | 26 | 4651 |
| ASN | 27 | 2856 | GLU | 3 | 3328 | GLU | 3 | 3658 | GLU | 3 | 3988 | ASN | 27 | 4429 |
| GLU | 3 | 2819 | ASN | 27 | 3274 | ASN | 27 | 3584 | ASN | 27 | 3957 | GLY | 25 | 4363 |
| LYS | 28 | 2470 | GLY | 25 | 2920 | GLY | 25 | 3358 | GLY | 25 | 3820 | GLU | 3 | 4314 |
| GLY | 25 | 2373 | LYS | 28 | 2842 | LYS | 28 | 3226 | LYS | 28 | 3618 | LYS | 28 | 4067 |
| ILE | 31 | 2113 | ILE | 31 | 2453 | ALA | 2 | 2712 | ALA | 30 | 3080 | ALA | 30 | 3506 |
| ALA | 2 | 1937 | ALA | 2 | 2382 | ILE | 31 | 2712 | ILE | 31 | 3029 | ILE | 31 | 3471 |
| ALA | 30 | 1898 | ALA | 30 | 2371 | ALA | 30 | 2696 | ALA | 2 | 3005 | ALA | 2 | 3358 |
| ILE | 32 | 1863 | GLY | 29 | 2255 | GLY | 29 | 2597 | GLY | 29 | 2921 | GLY | 29 | 3320 |
| GLY | 29 | 1809 | ILE | 32 | 2239 | ILE | 32 | 2524 | ILE | 32 | 2866 | ILE | 32 | 3261 |
| LEU | 34 | 1780 | LEU | 34 | 2104 | LEU | 34 | 2330 | LEU | 34 | 2582 | LEU | 34 | 2977 |
| ASP | 1 | 1531 | MET | 35 | 1837 | MET | 35 | 2104 | MET | 35 | 2408 | MET | 35 | 2792 |
| VAL | 36 | 1516 | VAL | 36 | 1818 | VAL | 36 | 2047 | VAL | 36 | 2296 | VAL | 36 | 2595 |
| MET | 35 | 1469 | ASP | 1 | 1815 | ASP | 1 | 2027 | ASP | 1 | 2251 | ASP | 1 | 2452 |
| VAL | 39 | 1350 | GLY | 33 | 1661 | GLY | 33 | 1883 | GLY | 33 | 2135 | GLY | 33 | 2451 |
| GLY | 33 | 1348 | VAL | 39 | 1629 | VAL | 39 | 1872 | VAL | 39 | 2115 | VAL | 39 | 2423 |
| ILE | 41 | 1208 | VAL | 40 | 1455 | ILE | 41 | 1675 | ILE | 41 | 1885 | GLY | 38 | 2126 |
| VAL | 40 | 1205 | ILE | 41 | 1454 | VAL | 40 | 1649 | VAL | 40 | 1848 | ILE | 41 | 2103 |
| GLY | 38 | 1060 | GLY | 37 | 1340 | GLY | 37 | 1571 | GLY | 38 | 1787 | VAL | 40 | 2094 |
| GLY | 37 | 1040 | GLY | 38 | 1336 | GLY | 38 | 1541 | GLY | 37 | 1781 | GLY | 37 | 2090 |
| ALA | 42 | 818 | ALA | 42 | 1006 | ALA | 42 | 1147 | ALA | 42 | 1299 | ALA | 42 | 1466 |

**Figure 3.**
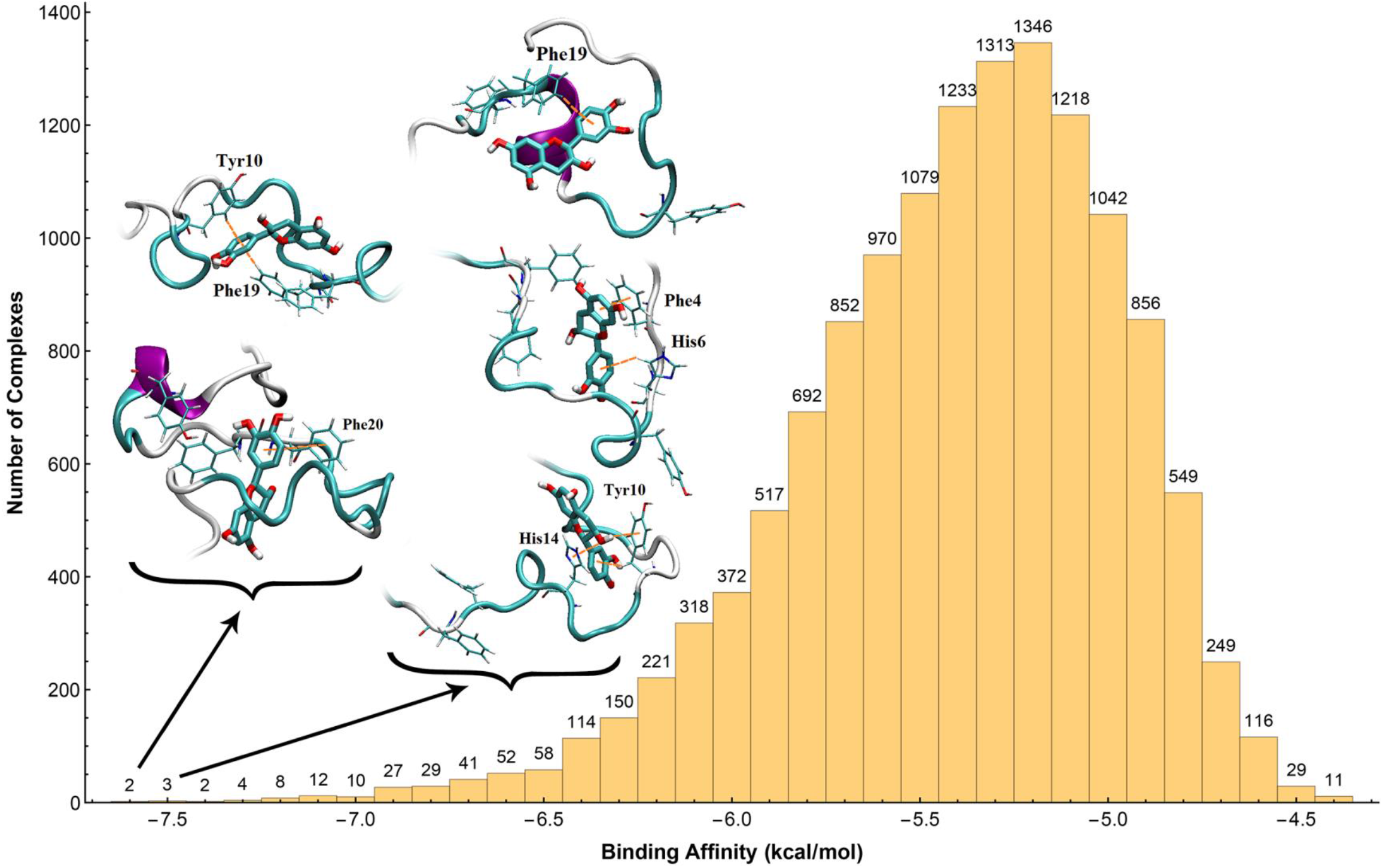
The histogram of binding affinity (kcal/mol) of the *C* ligand in set-1 (cutoff of 0.1 kcal/mol, Table 1). The binding affinities were divided to bins of 0.1 kcal/mol, and the numbers on top of the bins show the number of complexes in each bin. The three aromatic residue hotspots (*i.e*. Tyr10, Phe19, and Phe20, see Figure 1 for sequence) and interacting aromatic residues with the ligand for some complexes with the highest binding affinities are shown in licorice representation. Favorable aromatic interactions between the ligand and peptide are depicted with orange dashed lines.

**Figure 4.**
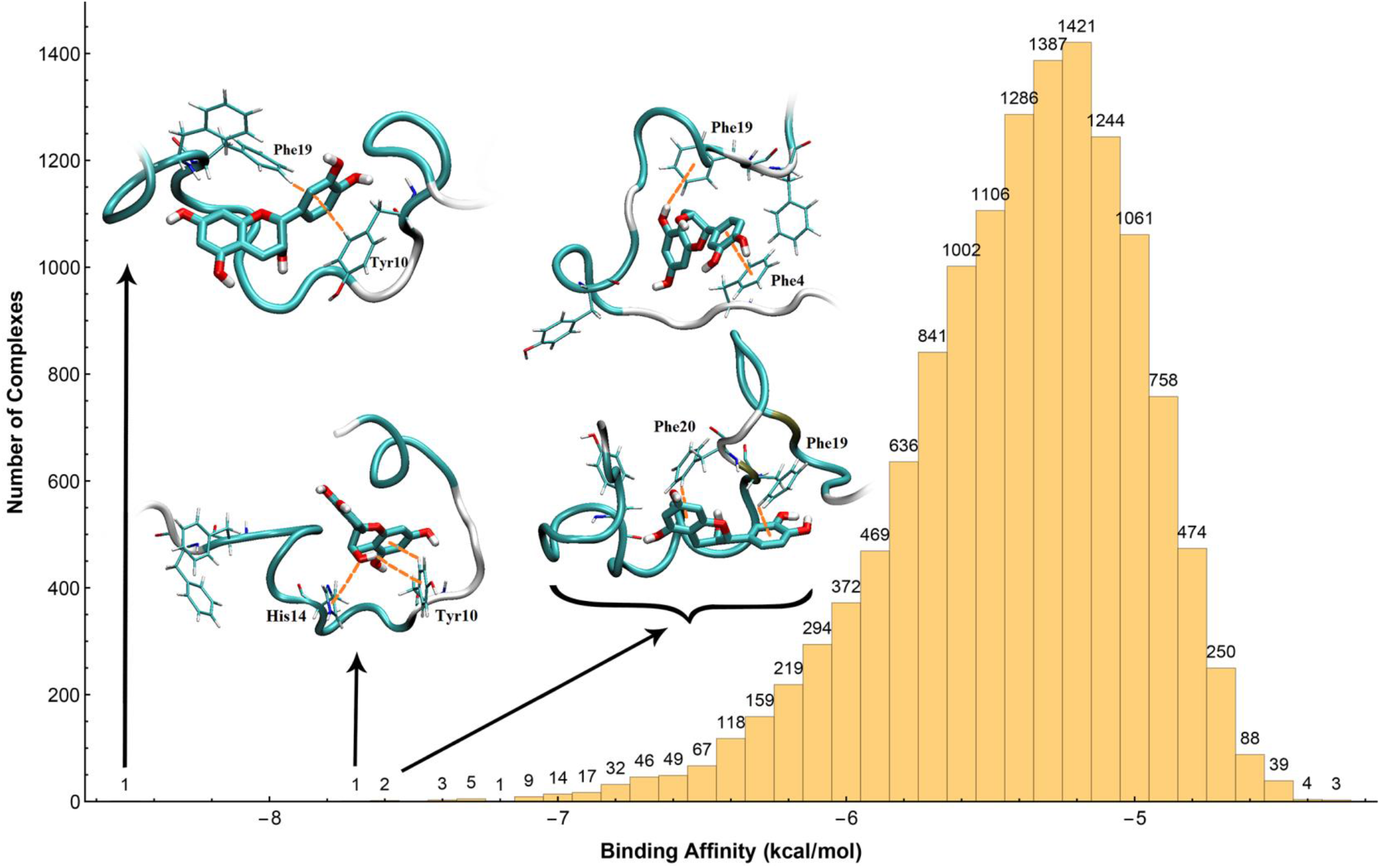
The histogram of binding affinity (kcal/mol) of the ***EC*** ligand over the set-1(cutoff of 0.1 kcal/mol, Table 1). The binding affinities were divided to bins of 0.1 kcal/mol, and the numbers on top of the bins show the number of complexes in each bin. The three aromatic residue hotspots (*i.e*. Tyr10, Phe19, and Phe20, see Figure 1 for sequence) and interacting aromatic residues with the ligand for some complexes with the highest binding affinities are shown in licorice representation. Favorable aromatic interactions between the ligand and peptide are depicted with orange dashed lines.

**Figure 5.**
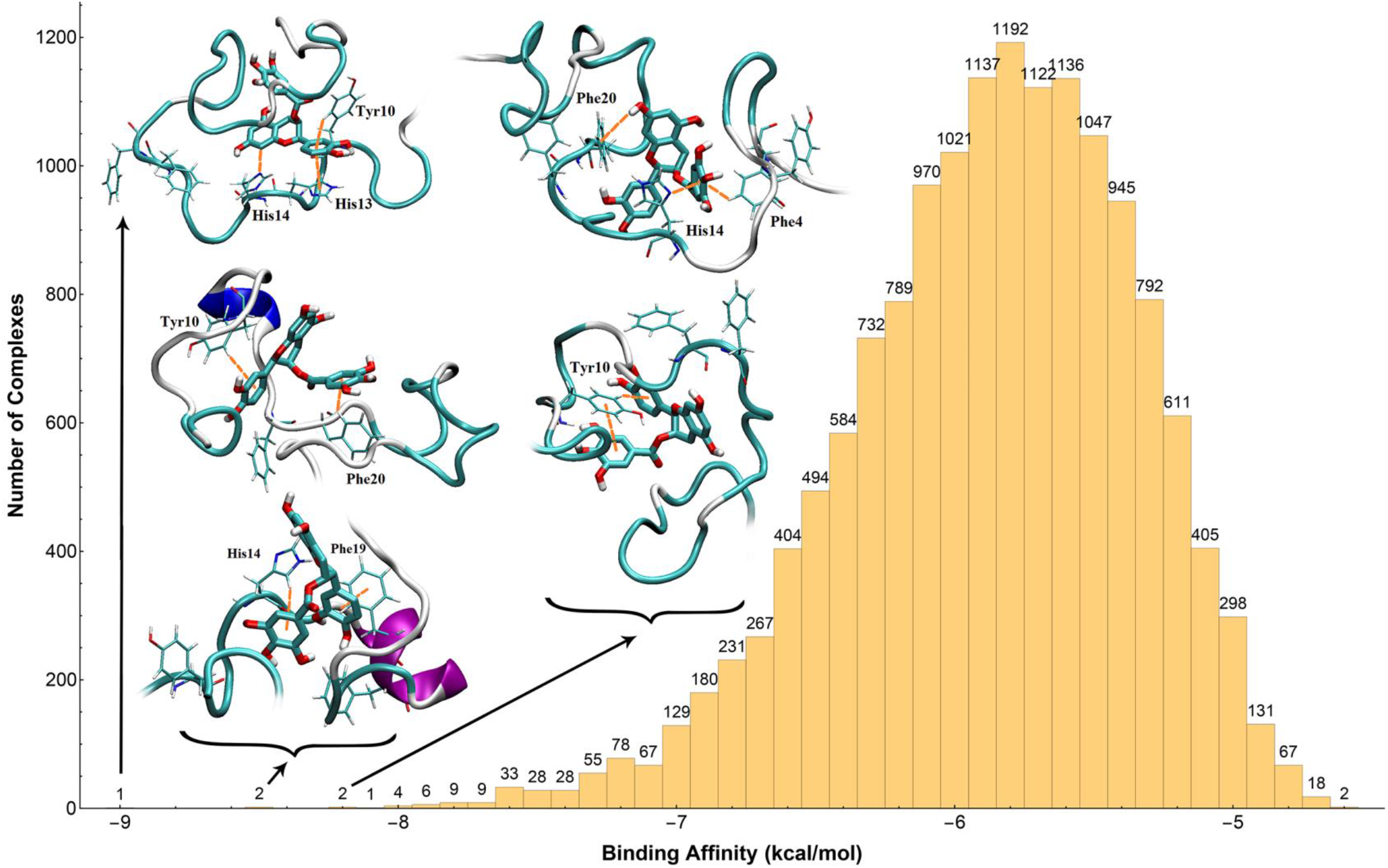
The histogram of binding affinity (kcal/mol) of the *ECG* ligand over the set-1 (cutoff of 0.1 kcal/mol, Table 1). The binding affinities were divided to bins of 0.1 kcal/mol, and the numbers on top of the bins show the number of complexes in each bin. The three aromatic residue hotspots (*i.e*. Tyr10, Phe19, and Phe20, see Figure 1 for sequence) and interacting aromatic residues with the ligand for some complexes with the highest binding affinities are shown in licorice representation. Favorable aromatic interactions between the ligand and peptide are depicted with orange dashed lines.

**Figure 6.**
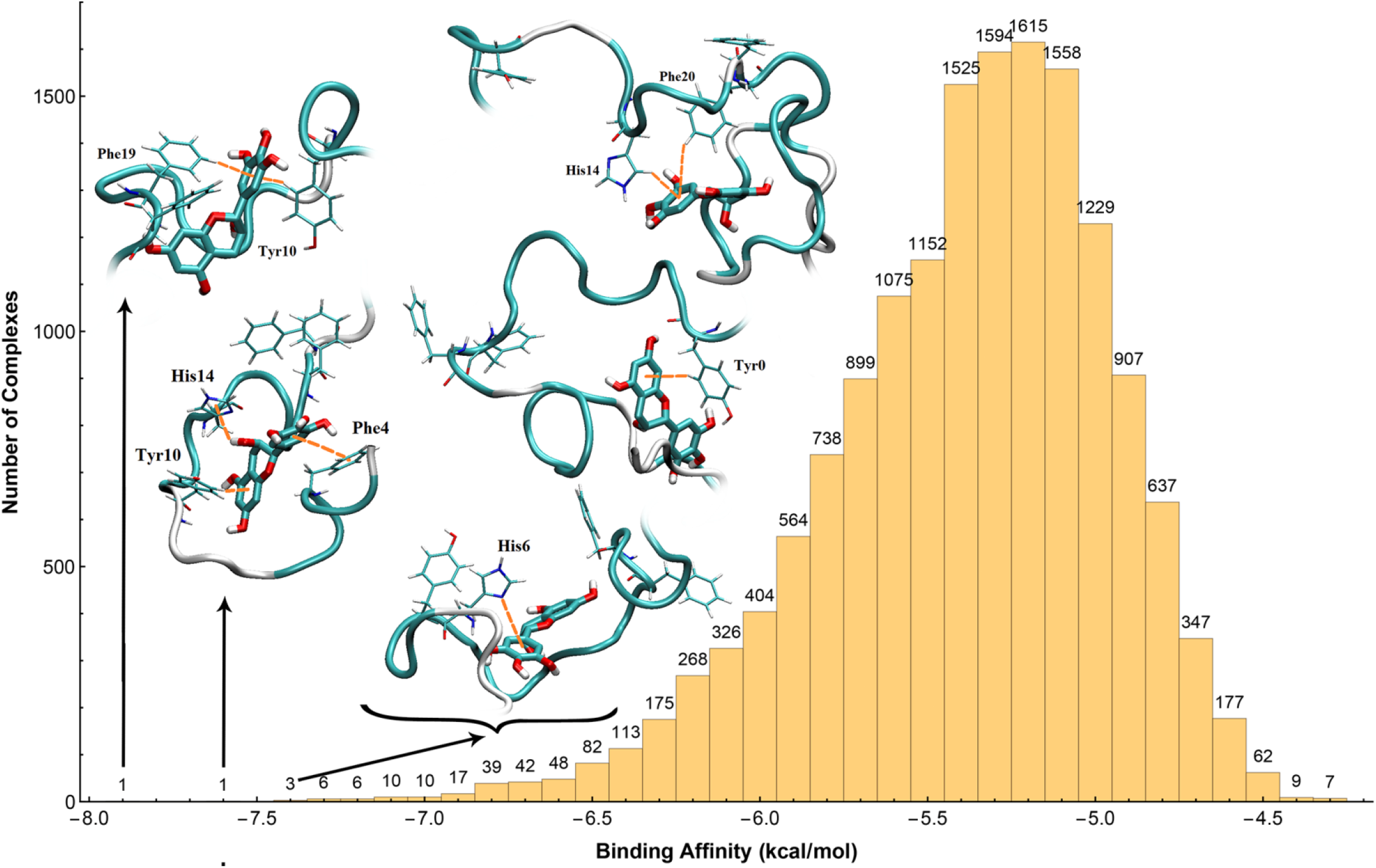
The histogram of binding affinity (kcal/mol) of the ***EGC*** ligand over the set-1 (cutoff of 0.1 kcal/mol, Table 1). The binding affinities were divided to bins of 0.1 kcal/mol, and the numbers on top of the bins show the number of complexes in each bin. The three aromatic residue hotspots (*i.e*. Tyr10, Phe19, and Phe20, see Figure 1 for sequence) and interacting aromatic residues with the ligand for some complexes with the highest binding affinities are shown in licorice representation. Favorable aromatic interactions between the ligand and peptide are depicted with orange dashed lines.

**Figure 7.**
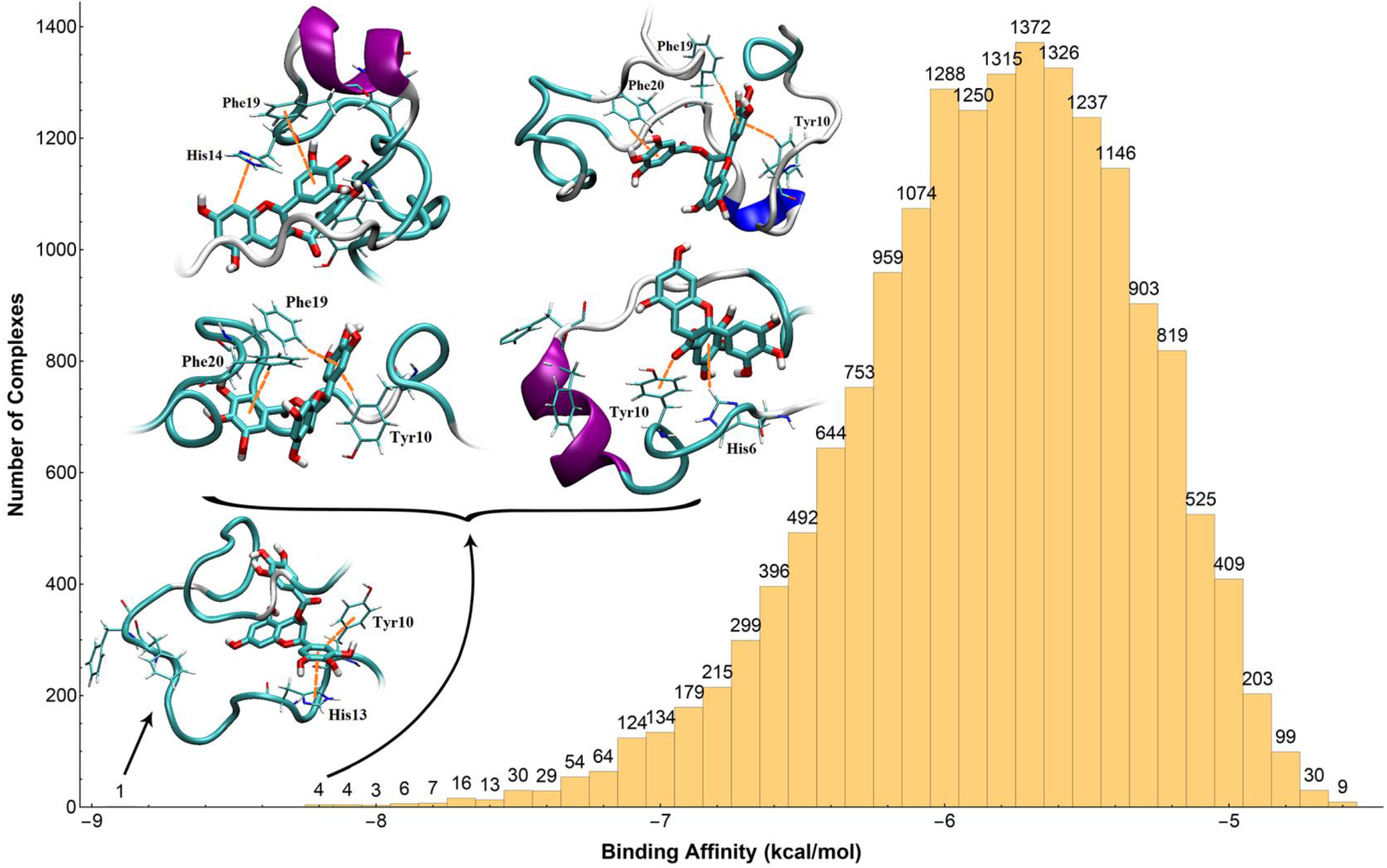
The histogram of binding affinity (kcal/mol) of the *EGCG* ligand over the set-1 (cutoff of 0.1 kcal/mol, Table 1). The binding affinities were divided to bins of 0.1 kcal/mol, and the numbers on top of the bins show the number of complexes for each bin. The three aromatic residue hotspots (*i.e*. Tyr10, Phe19, and Phe20, see Figure 1 for sequence) and interacting aromatic residues with the ligand for some complexes with the highest binding affinities are shown in licorice representation. Favorable aromatic interactions between the ligand and peptide are depicted with orange dashed lines.

The distributions of binding affinities (kcal/mol) of all ligands in set-1 (cutoff of 0.1 kcal/mol) are shown in **Figures 3-7**. In addition, the figures show a couple of complexes with the highest binding affinities and for each complex the different types of aromatic interactions, such as π-π, XH-π (X = C, N, O), and lone pair-π interactions between the aromatic rings of the ligands, and the sidechain of aromatic residues have been highlighted by colored dashed-lines for clarity. In all cases, at least one aromatic residue was in contact with the ligand and in most cases, catechin compounds possessing multiple aromatic rings were capable of interacting with several aromatic residues simultaneously. Therefore, it seems that these aromatic interactions play an important role in binding to aggregation-prone regions of A*β*_42_ and are essential for high affinity and binding specificity.

### 3.2. Analysis of MD Simulations

In order to evaluate the docking results, microsecond-scale MD simulations were performed on the A*β*_42_-L (L = *C*, *EC*, *ECG, EGC, EGCG*) structures for which the largest binding energies were obtained by the docking procedure. The initial and final MD structures are provided in **Figure S1** which show that the ligands maintain their interactions with A*β*_42_ throughout the simulation. Root mean square deviation (RMSD) of the A*β*_42_ indicates that in all cases the system has reached steady state (**Table S1** and **Figure S2**). The stability of the A*β*_42_-L complexes is also reflected in the relatively low radius of gyration (~1.0 nm) of the A*β*_42_ (**Table S1** and **Figure S3**). Moreover, the steady solvent accessible surface area (SASA; with < 7% fluctuations) observed for the chains provides further support for the stability and compactness of the system (**Table S1** and **Figure S4**).

The docking results suggest that Tyr10, Phe19, and to some extent Phe20, are the main residues involved in ligand binding. Average distances between these residues and the five ligands from the MD simulations are provided in **Table 7**. In agreement with the results from our docking procedure, with the exception of *C* for which the distance is ~1.0 nm, the average distance between the ligands and all three residues is ~0.5 nm (**Table 7** and **Figures S5 - S9**). The time evolution of these distances and the end-to-end distance for A*β*_42_ are shown in **Figures S5 - S9**.

**Table 7.** Average A*β*_42_ end-to-end distance (D1-A42) and distances between EGCG and Y10, F19, F20 residues (the amino acid sequence is provided in the caption of Figure 1). Standard deviations are provided in parentheses.

|  | Distance (nm) |  |  |  |  |
| --- | --- | --- | --- | --- | --- |
|  | C | EC | ECG | EGC | EGCG |
| **end-to-end** | 1.10 (0.66) | 1.69 (0.73) | 1.30 (0.48) | 1.77 (0.60) | 1.54 (0.59) |
| **Ligand...Y10** | 1.08 (0.45) | 0.47 (0.40) | 0.54 (0.18) | 0.29 (0.08) | 0.70 (0.22) |
| **Ligand...F19** | 0.97 (0.54) | 0.95 (0.51) | 0.55 (0.25) | 0.50 (0.20) | 0.40 (0.23) |
| **Ligand...F20** | 0.74 (0.53) | 0.72 (0.39) | 0.44 (0.24) | 0.44 (0.20) | 0.36 (0.26) |

Inspection of hydrogen bonds between the ligands and A*β*_42_ reveals that hydrogen bonding plays a more important role in binding of *EGCG* and *EGC* than with the other three ligands (**Table 8**). Binding of *C* seems to have the least dependence on the hydrogen bonding among the ligands and relies on the π–π, XH–π (X = C, O) interactions. Average number of A*β*_42_···ligand, A*β*_42_…solvent, and ligand···solvent hydrogen bonds are collected in **Table 8**.

**Table 8.** Average number of hydrogen bonds (donor···acceptor ≤ 0.35 nm and α (∠(Hydrogen-Donor-Acceptor) ≤ 30°). Standard deviations are provided in parentheses.

|  | Number of hydrogen bonds (SD) |  |  |  |  |
| --- | --- | --- | --- | --- | --- |
|  | C | EC | ECG | EGC | EGCG |
| **$A\beta$...Ligand** | 2 (2) | 2 (1) | 4 (2) | 4 (1) | 5 (2) |
| **$A\beta$...Solvent** | 109 (17) | 114 (20) | 108 (21) | 114 (21) | 112 (8) |
| **Ligand...Solvent** | 6 (2) | 5 (3) | 7 (4) | 5 (3) | 7 (3) |

**Figures S10 - S14** depict the contributions of A*β*_42_ residues that involve hydrogen bonding with the ligands. A comparison among the five A*β*_42_···ligand systems reveals that there are a few residues in the A*β*_42_ sequence that frequently form hydrogen bonds with the five different catechins (e.g., Glu22, Asp23, Ala42, etc.). These intransient hydrogen bonds alternate between the ligand and the residues, which may suggest that hydrogen bonds have a less important role in the binding of the ligands to A*β*_42_ compared to stacking interactions. The average number of hydrogen bonds between all the amino acids of the peptide and ligands are depicted in **Figure S15**. Similar conclusion has been previously made for A*β*_42_ protofibrils and *EGCG*, where *EGCG* was shown to bind to A*β*_42_ monomer through hydrophobic, π–π stacking, and hydrogen bonds.^120^ Moreover, same study identifies Asp1, Glu22, and Ala42 residues to form the most hydrogen bonds with *EGCG,* which except for Asp1, agrees with our observation (**Figure S14**). It should be noted, however, that while we have examined disordered structures in this study, Li *et al*.^120^ considered fibrils and 25 *EGCG* molecules.

## 4. Conclusions

In this work, binding of various well-known catechins present in green tea to the amyloid-*β* peptide (A*β*) has been predicted and analyzed. For this purpose, a computational pipeline in the framework of the ensemble docking strategy has been proposed in which a structurally heterogeneous ensemble of conformations of A*β*_42_ is used. The ensemble is generated by the Blockwise Excursion Sampling (BES) protocol^70^ in which the conformational sampling is performed on the basis of many uncorrelated short-time MD simulations starting from different reasonable points of the accessible phase space.

It was observed that all green tea catechins compounds tended to interact with the aromatic residues through stacking and/or T-shaped interactions and, because of this, all compounds show a high tendency to interact with the hydrophobic region of A*β*_42_ spanning residues from Tyr10 to Phe20, the region with the highest number of the aromatic residues in full-length A*β*_42_. This region also encompasses the central hydrophobic core (CHC, residues 16 - 20) that, based on many experimental and computational studies, plays a key role in the aggregation process of A*β*_42_. Therefore, the docking results indicate that all studied ligands, especially the *EGCG*, can act as potent inhibitors against amyloid aggregation by blocking the central hydrophobic core. Additionally, it has been suggested that both hydrophobic aromatic interactions and hydrogen bonding are crucial for the binding of catechins to A*β*_42_.

To evaluate the obtained findings in binding of catechin compounds to A*β*_42_, long multimicrosecond MD simulations were performed. It was shown that the present docking protocol is highly successful in identifying catechins’ binding pockets in monomeric A*β*_42_, in agreement with previous MD simulations and some recent experimental observations for similar A*β*_42_-catechin complexes.^28, 71–73, 77–79^ Finally, we suggested that our proposed pipeline with low computational cost in comparison with MD simulations is a suitable approach for high-throughput structure-based virtual screening of ligand libraries against the intrinsically disordered proteins (IDPs), such as the A*β*.

## Data and Software Availability

GAMESS package and AutoDock Vina (version 1.1.2) were used under a free academic license for ligands preparation and docking simulations. Produced and analyzed data are available upon request. MD simulations were performed using GROMACS 2016.3.

## Supporting Information

RMSD, R_g_, and SASA of A*β*_42_ in the different systems: Table S1 (standard deviations in parentheses). Initial and final structures of A*β*_42_ in the presence of ligands: Figure S1. RMSD, radius of gyration and solvent accessible surface area of the A*β*_42_ backbone in the presence of different ligands, Figures S2, S3 and S4, respectively. Figures S5-S9 show the average A*β*_42_ end-to-end distance and distances between residues Y10, F19 and F20, and *C, EC, ECG, EGC* and *EGCG*, respectively. Figures S10-S14: Hydrogen bonds between selected residues of **A*β*_42_** and *C, EC, ECG, EGC* and *EGCG,* respectively. Figure S15: Average number of hydrogen bonds between A*β*_42_ amino acids and the ligands.

## Author Contributions

All authors contributed to the design, implementation and analysis of the research, interpretation of the results, and writing of the manuscript. All authors have given their approval for the final version of the manuscript.

## Acknowledgments

C.C.G is recipient of an Ontario Trillium Scholarship (OTS). M.K. thanks NSERC Canada Research Chairs Program the Natural Sciences and Engineering Research Council of Canada (NSERC) for financial support. Computational resources were provided by SharcNet and Compute Canada.

## Supporting Information

**Table S1.** RMSD, R_g_, and SASA of A*β*_42_ in the different systems (standard deviations in parentheses).

|  | Systems |  |  |  |  |
| --- | --- | --- | --- | --- | --- |
|  | C | EC | ECG | EGC | EGCG |
| **RMSD (nm)** | 0.85 (0.12) | 1.26 (0.09) | 0.7 (0.09) | 1.26 (0.17) | 0.99 (0.14) |
| **R<sub>g</sub> (nm)** | 0.96 (0.09) | 0.98 (0.07) | 0.96 (0.07) | 0.99 (0.07) | 1.01 (0.06) |
| **SASA (nm<sup>2</sup>)** | 35.56 (3.19) | 36.82 (2.53) | 36.96 (2.03) | 37.07 (1.91) | 38.19 (1.89) |

**Figure S1.**
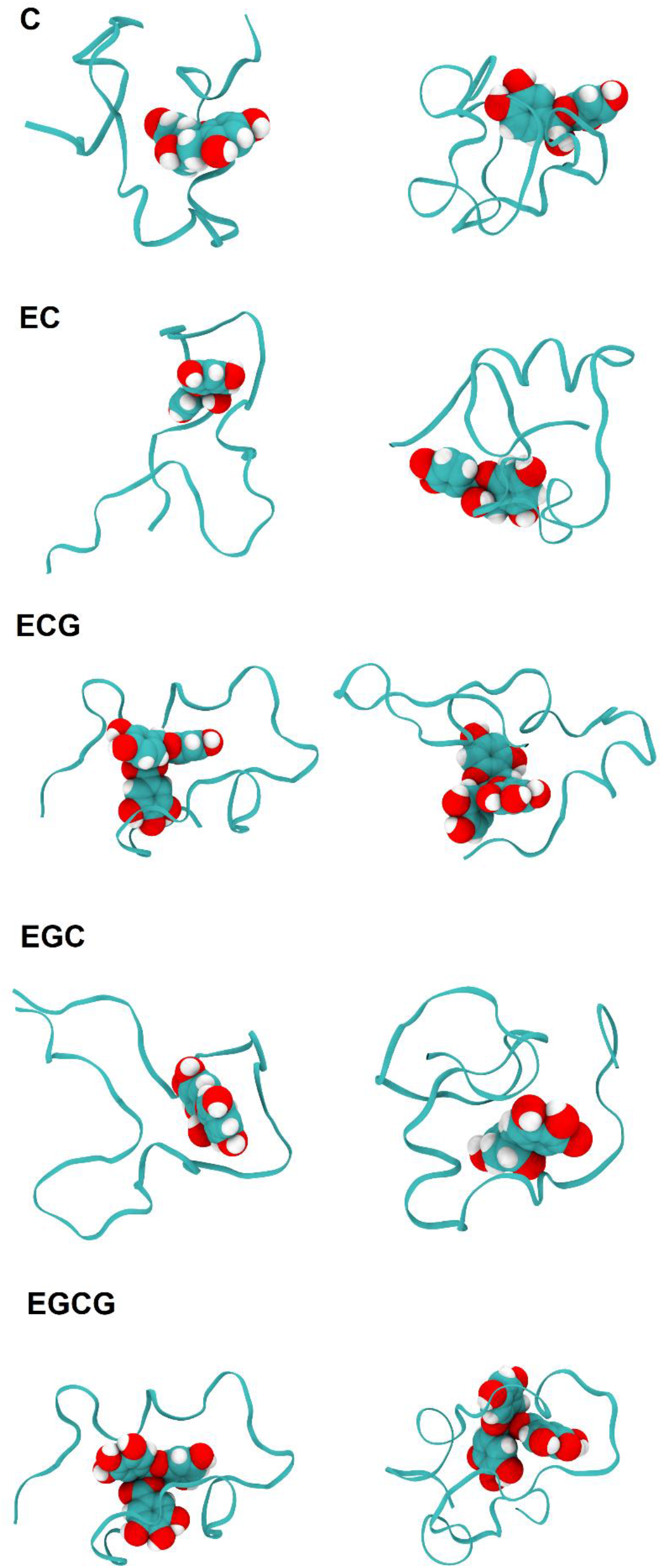
Initial (left) and final (right, at 3.0 μs) structures of A*β*_42_ in the presence of ligands. A*β*_42_ is shown using ribbons and the ligands using the van der Waals representation.

**Figure S2.**
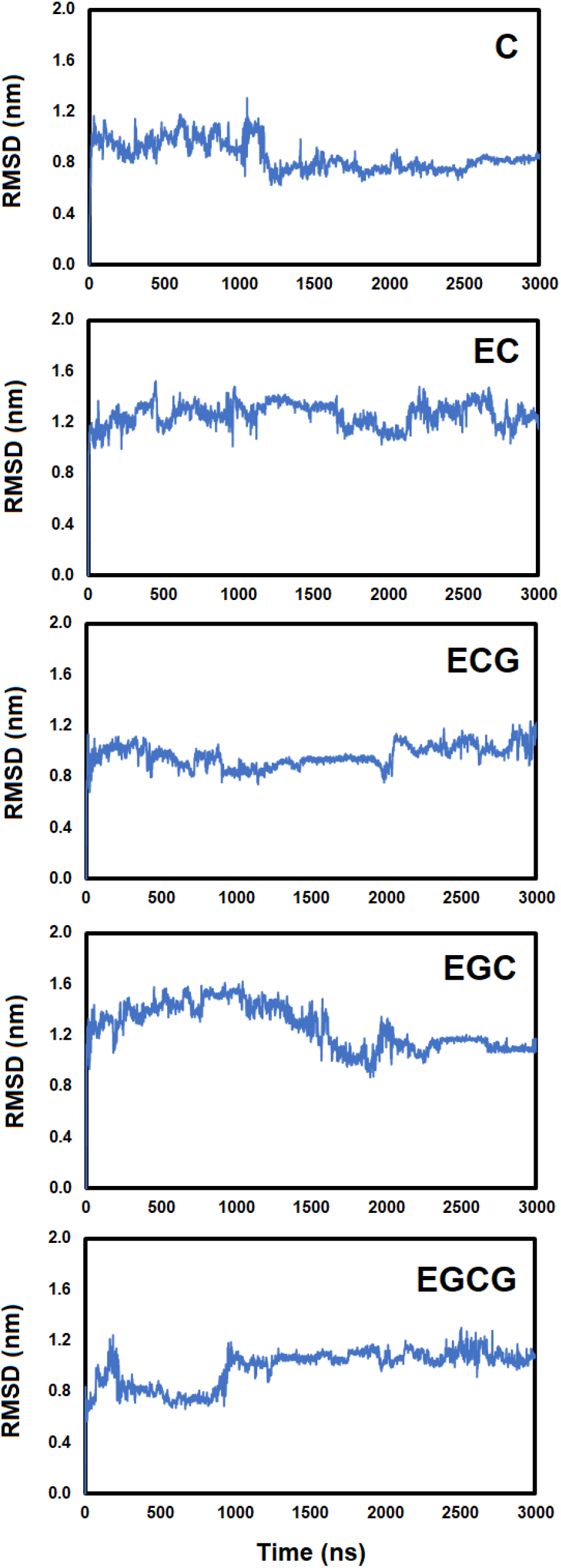
RMSD of A*β*_42_ backbone in the presence of different ligands.

**Figure S3.**
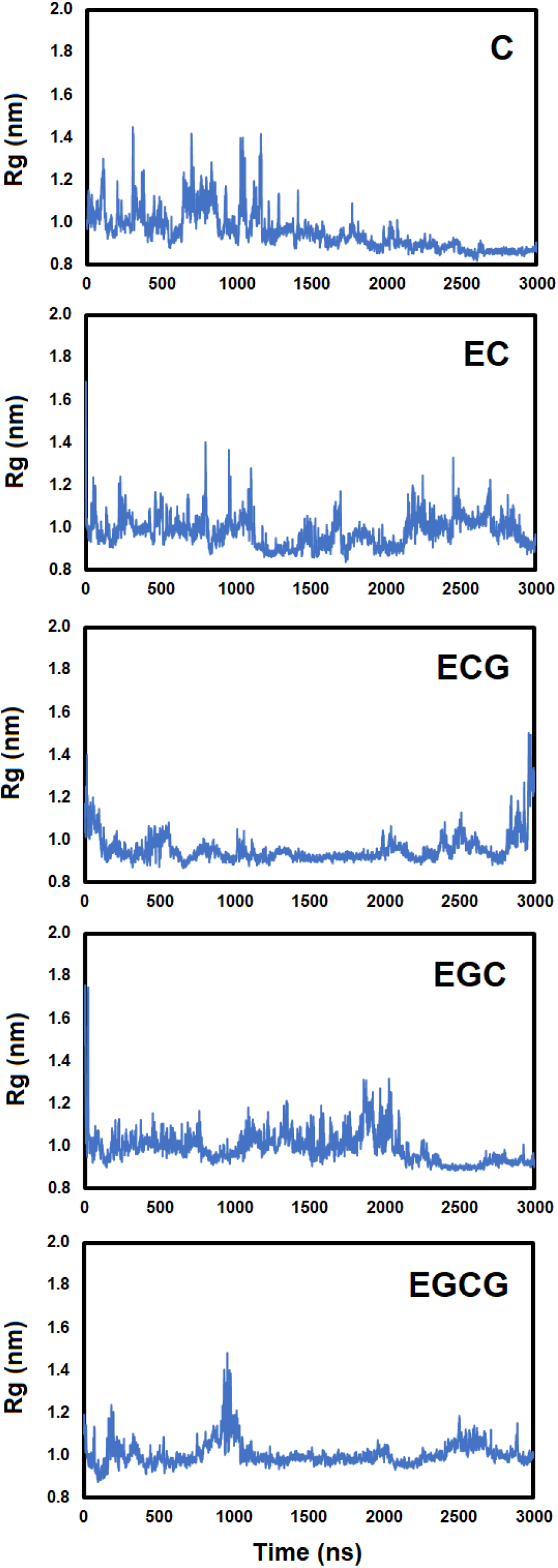
Radius of gyration of A*β*_42_ backbone in the presence of different ligands.

**Figure S4.**
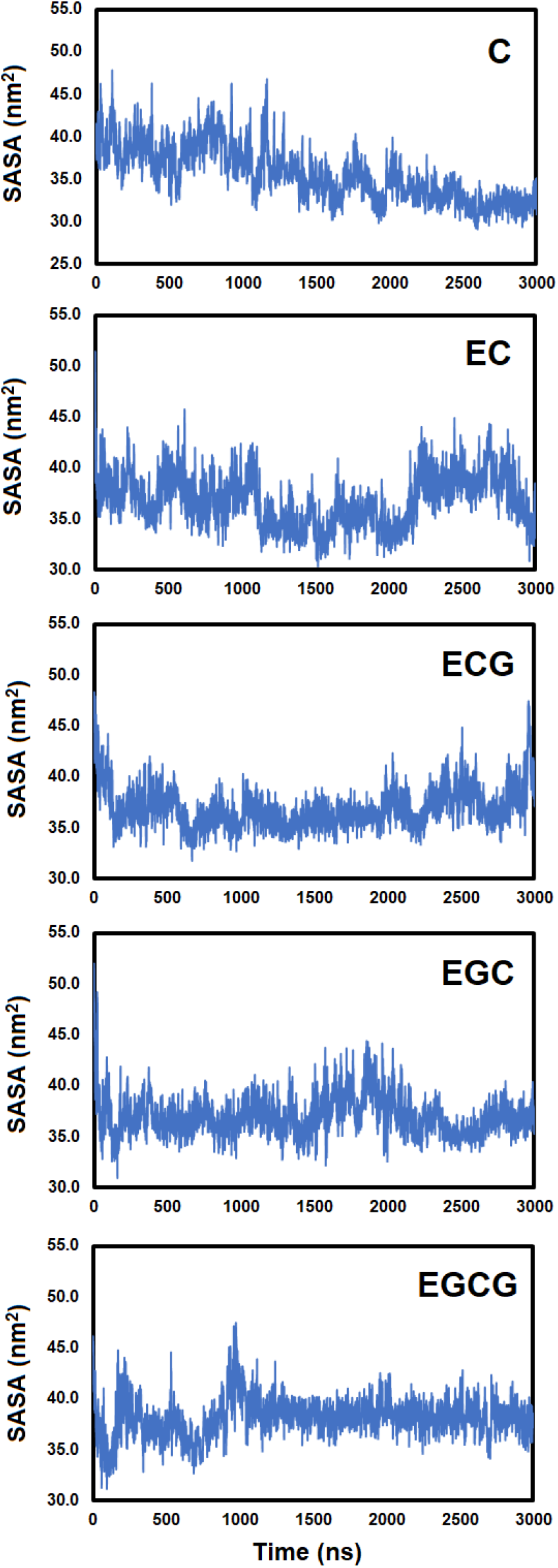
Solvent accessible surface area of A*β*_42_ backbone in the presence of different ligands. SASA was calculated based on the numerical Double Cubic Lattice Method (DCLM)^1^ as implemented in the GROMACS package using 1 ns intervals.

**Figure S5.**
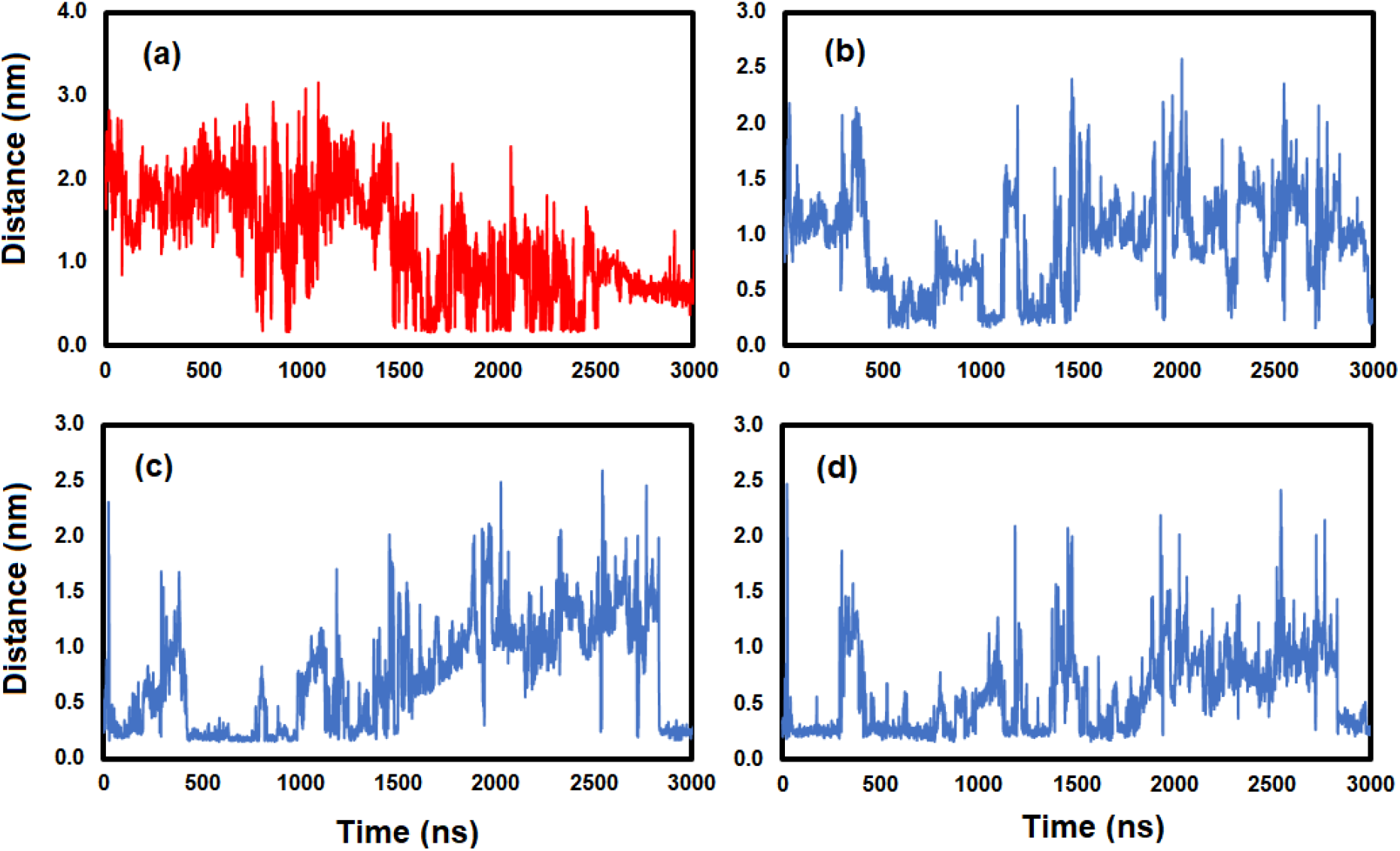
(a) Average A*β*_42_ end-to-end distance (D1-A42) and distances between **C** and (b) Y10, (c) F19, and (d) F20 residues.

**Figure S6.**
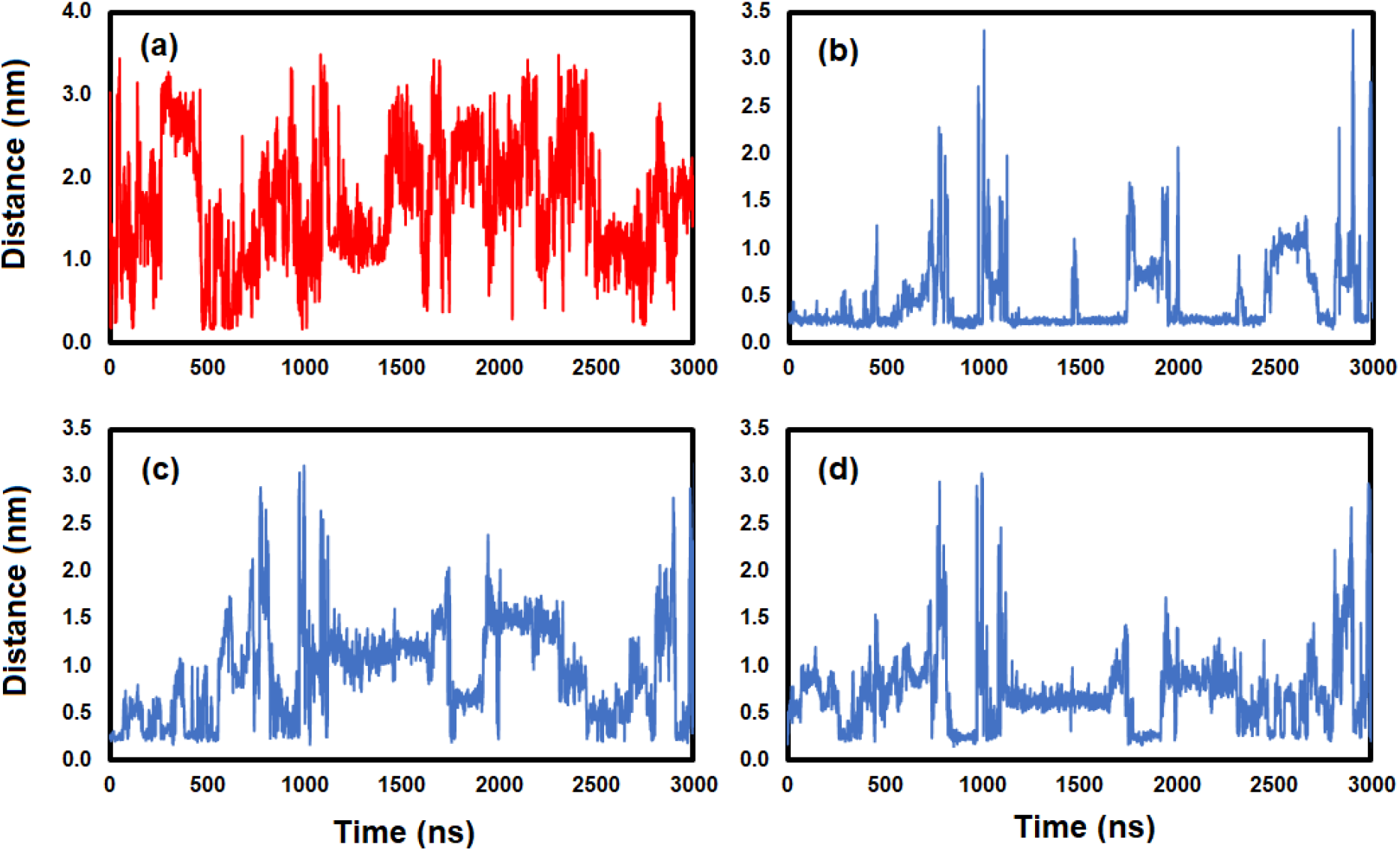
(a) Average A*β*_42_ end-to-end distance (D1-A42) and distances between **EC** and (b) Y10, (c) F19, and (d) F20 residues.

**Figure S7.**
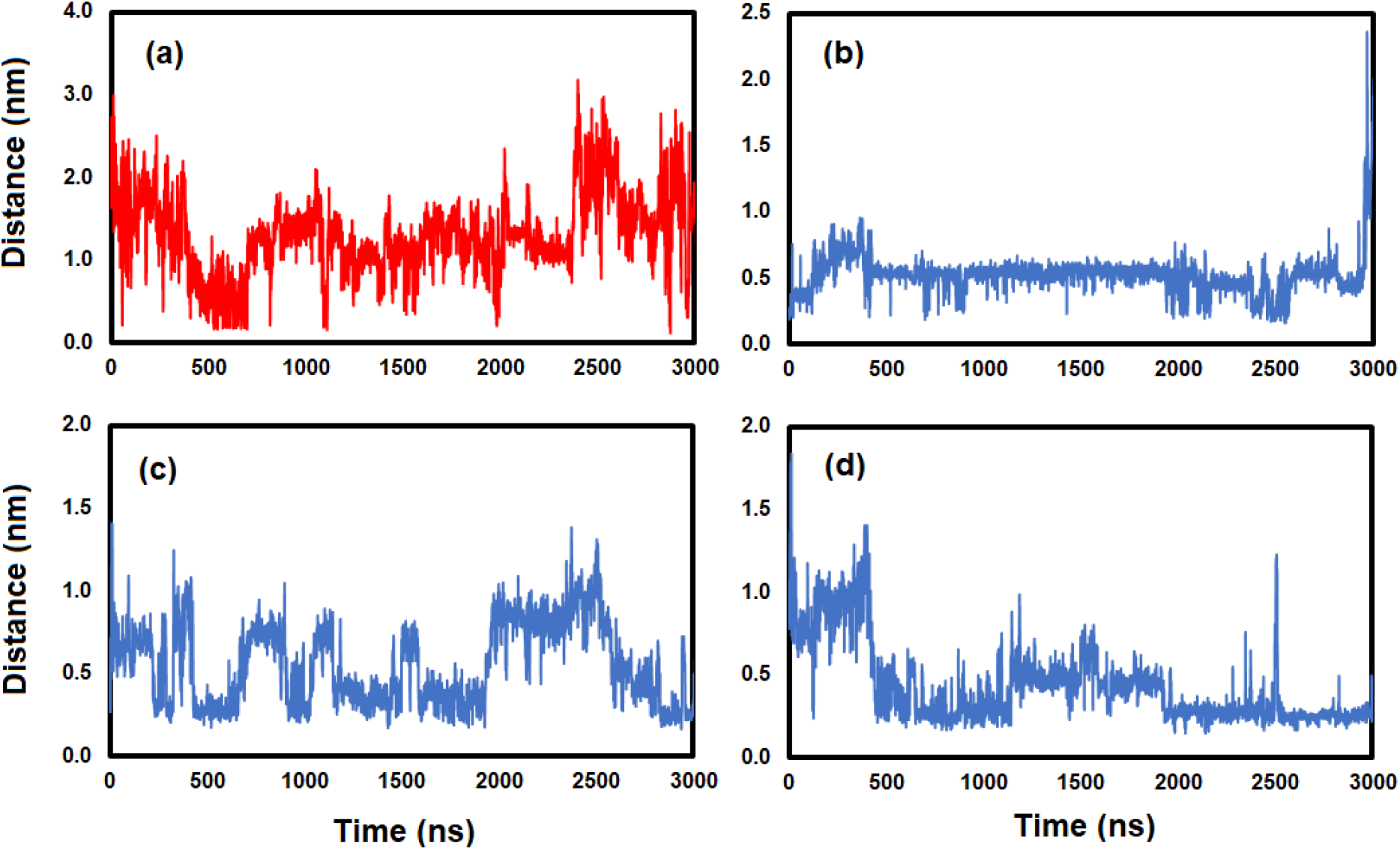
(a) Average A*β*_42_ end-to-end distance (D1-A42) and distances between ECG and (b) Y10, (c) F19, and (d) F20 residues.

**Figure S8.**
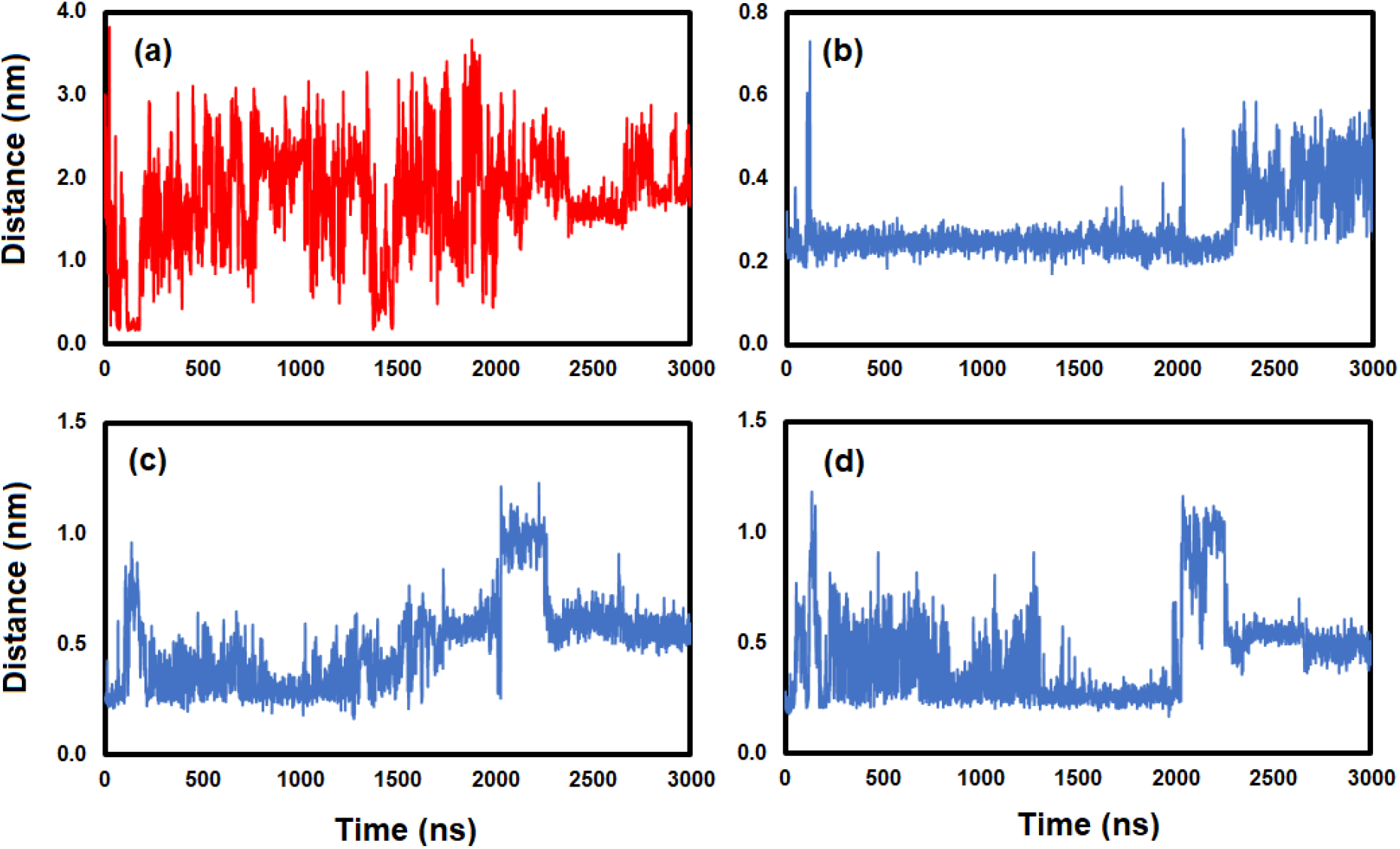
(a) Average A*β*_42_ end-to-end distance (D1-A42) and distances between EGC and (b) Y10, (c) F19, and (d) F20 residues.

**Figure S9.**
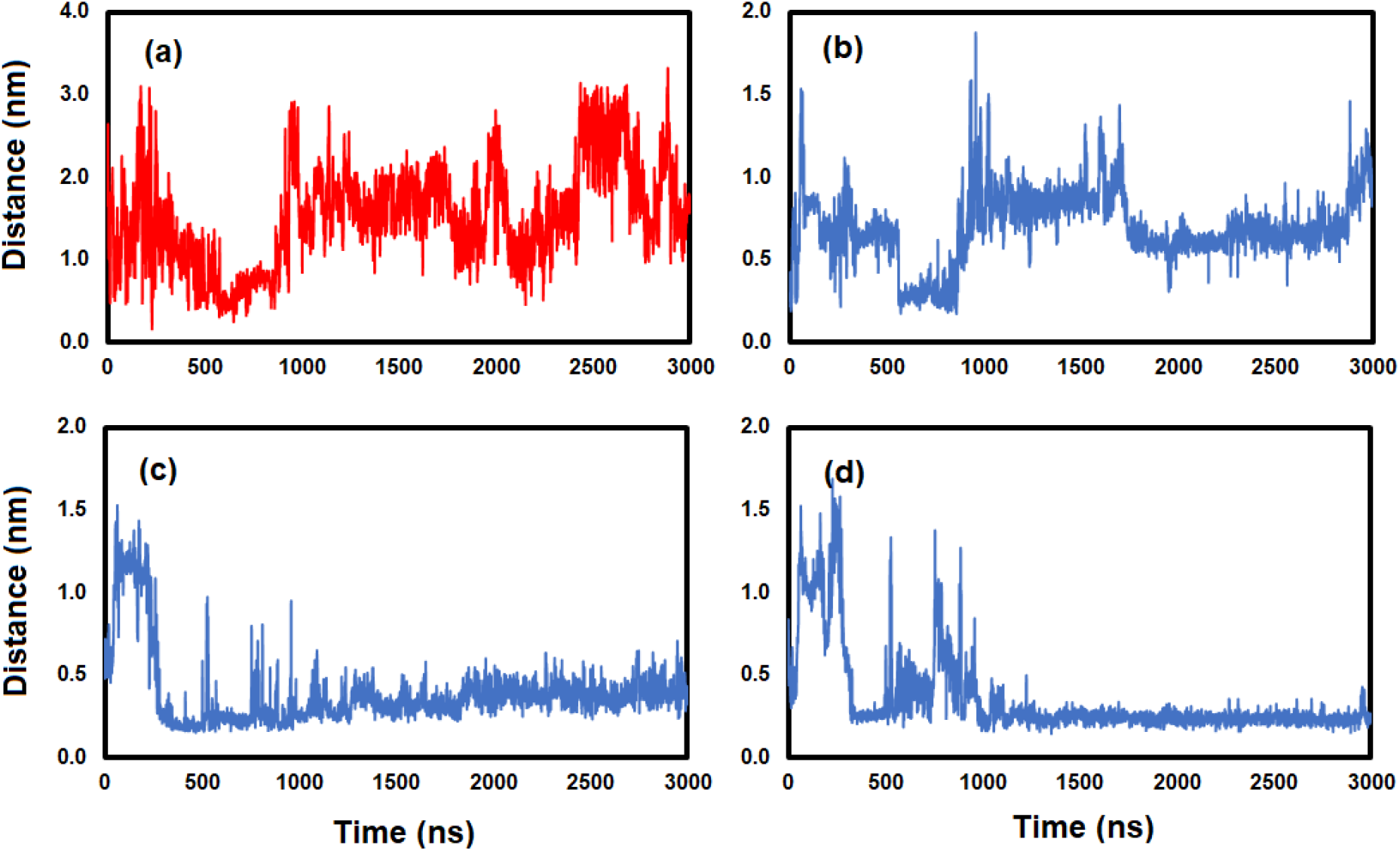
(a) Average A*β*_42_ end-to-end distance (D1-A42) and distances between EGCG and (b) Y10, (c) F19, and (d) F20 residues.

**Figure S10.**
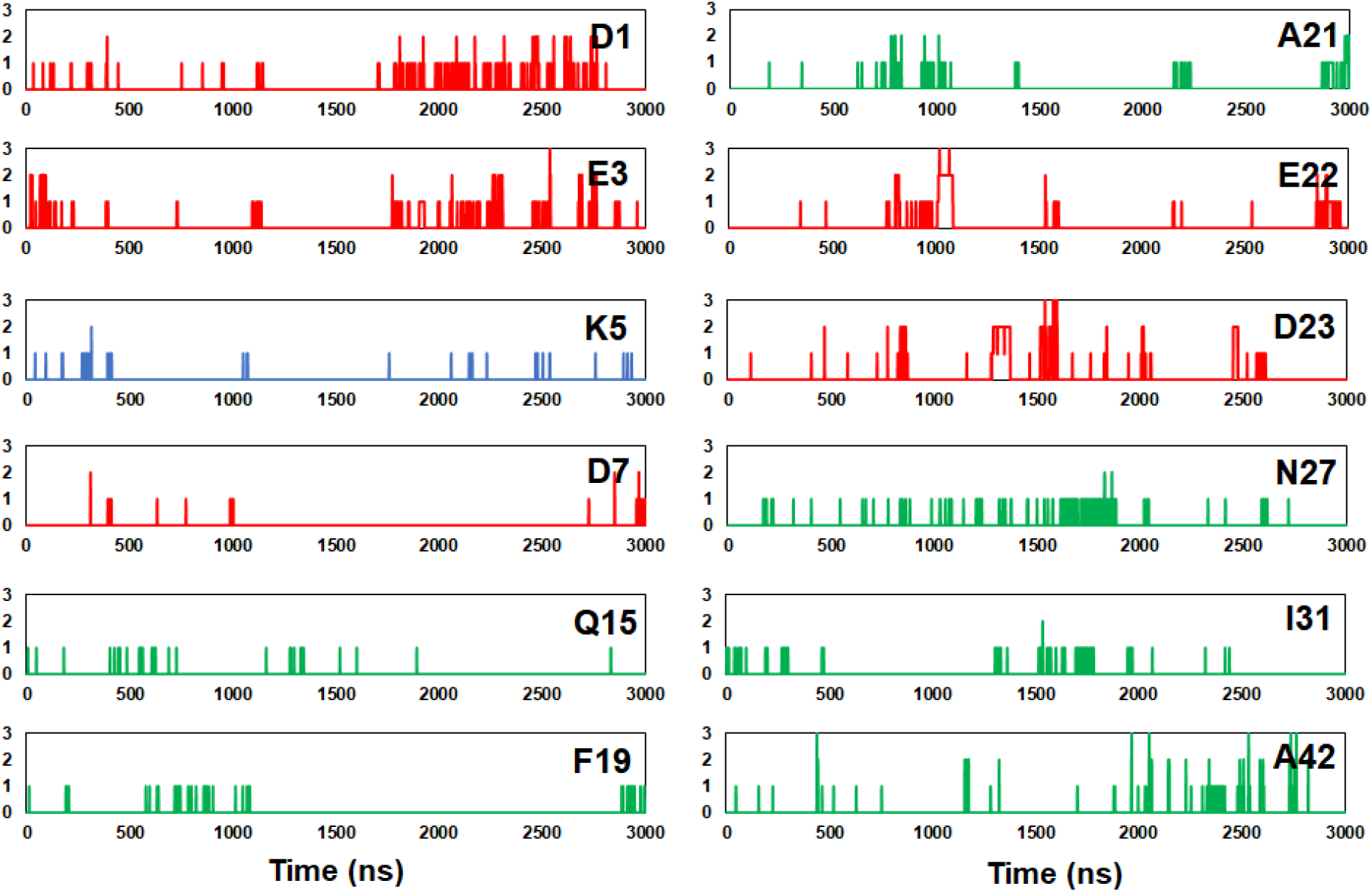
Hydrogen bonds between C and A*β*_42_ selected residues. Anionic, cationic, and neutral residues are shown in red, blue, and green, respectively.

**Figure S11.**
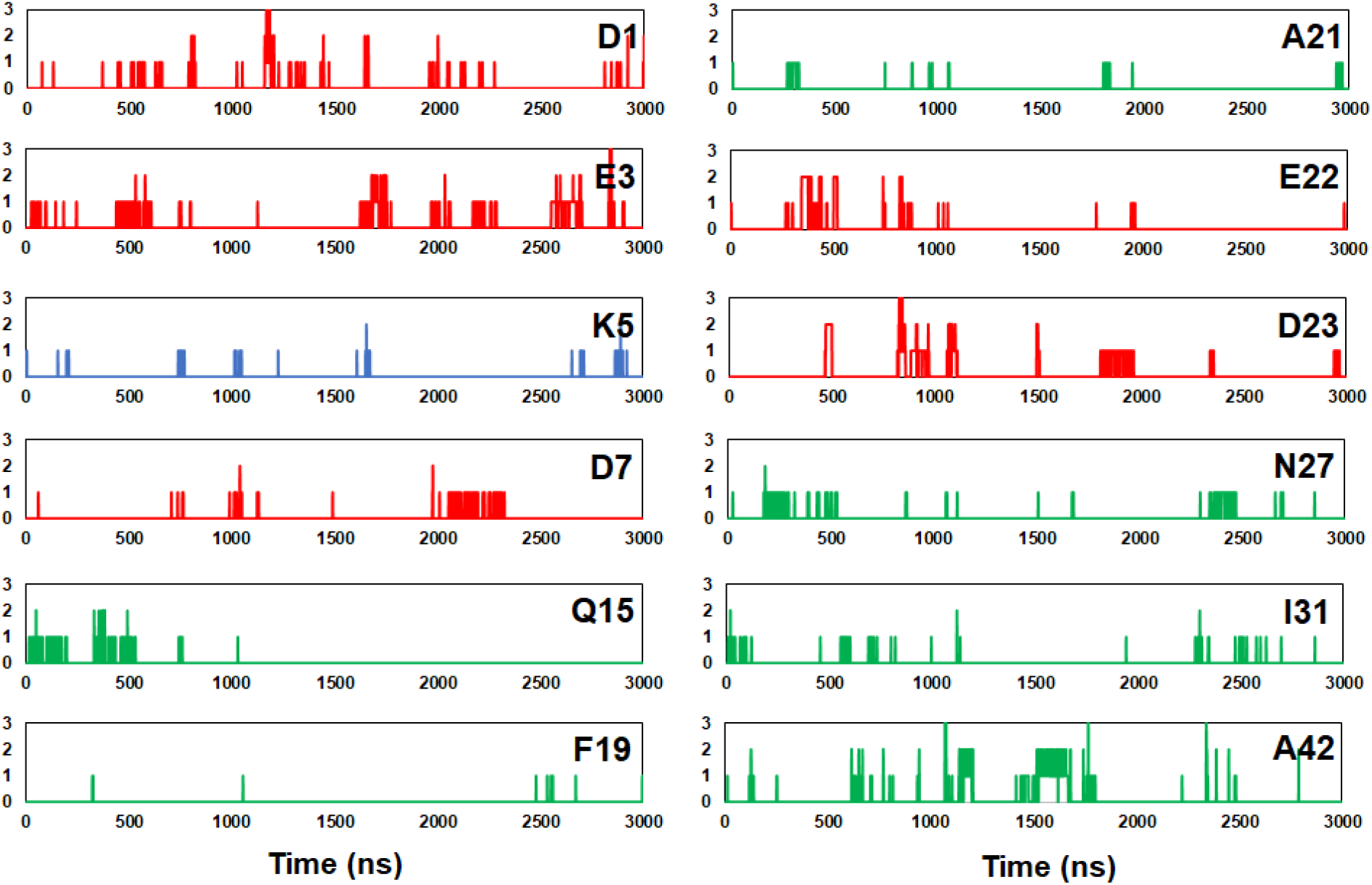
Hydrogen bonds between EC and A*β*_42_ selected residues. Anionic, cationic, and neutral residues are shown in red, blue, and green, respectively.

**Figure S12.**
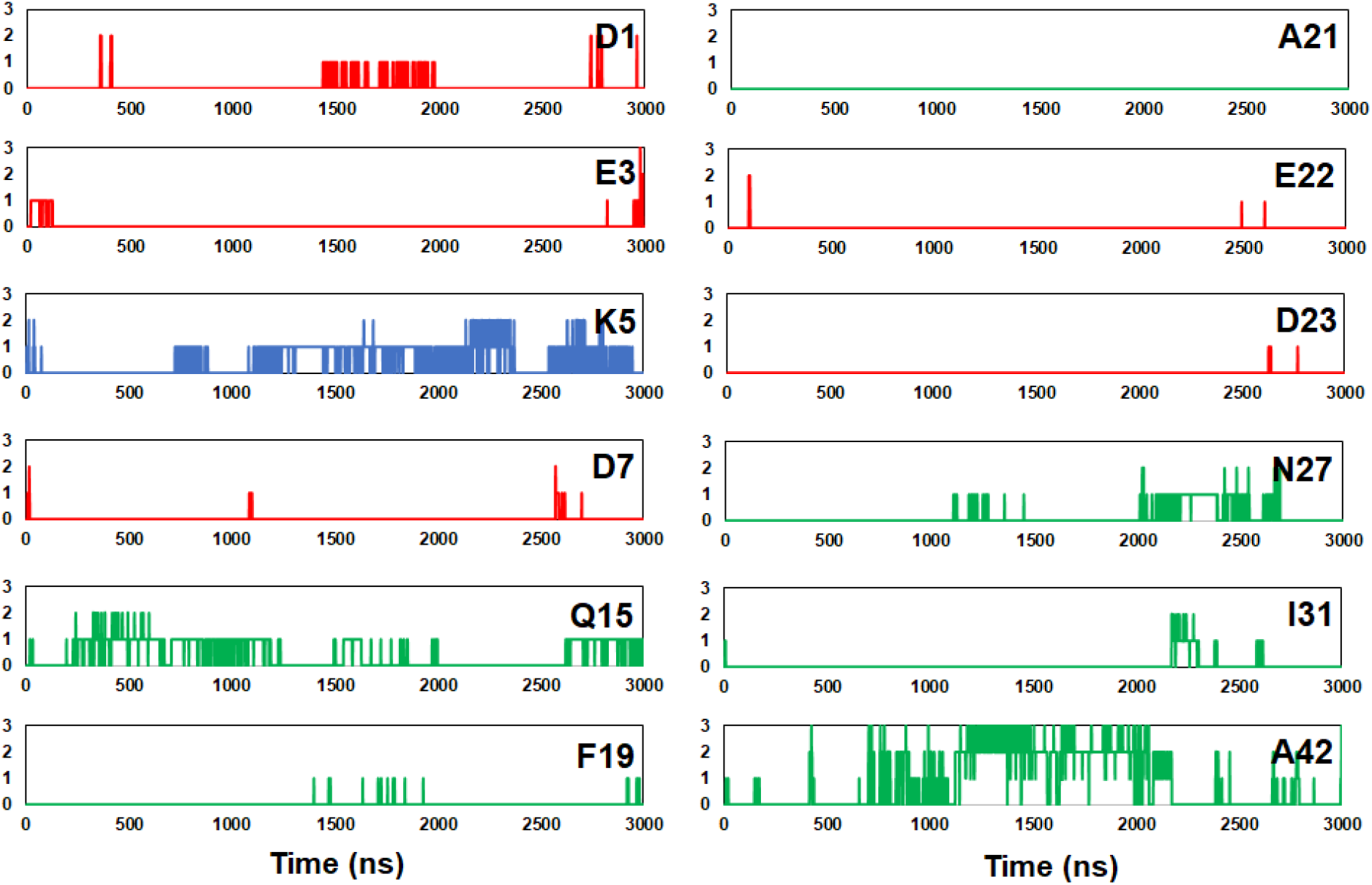
Hydrogen bonds between ECG and A*β*_42_ selected residues. Anionic, cationic, and neutral residues are shown in red, blue, and green, respectively.

**Figure S13.**
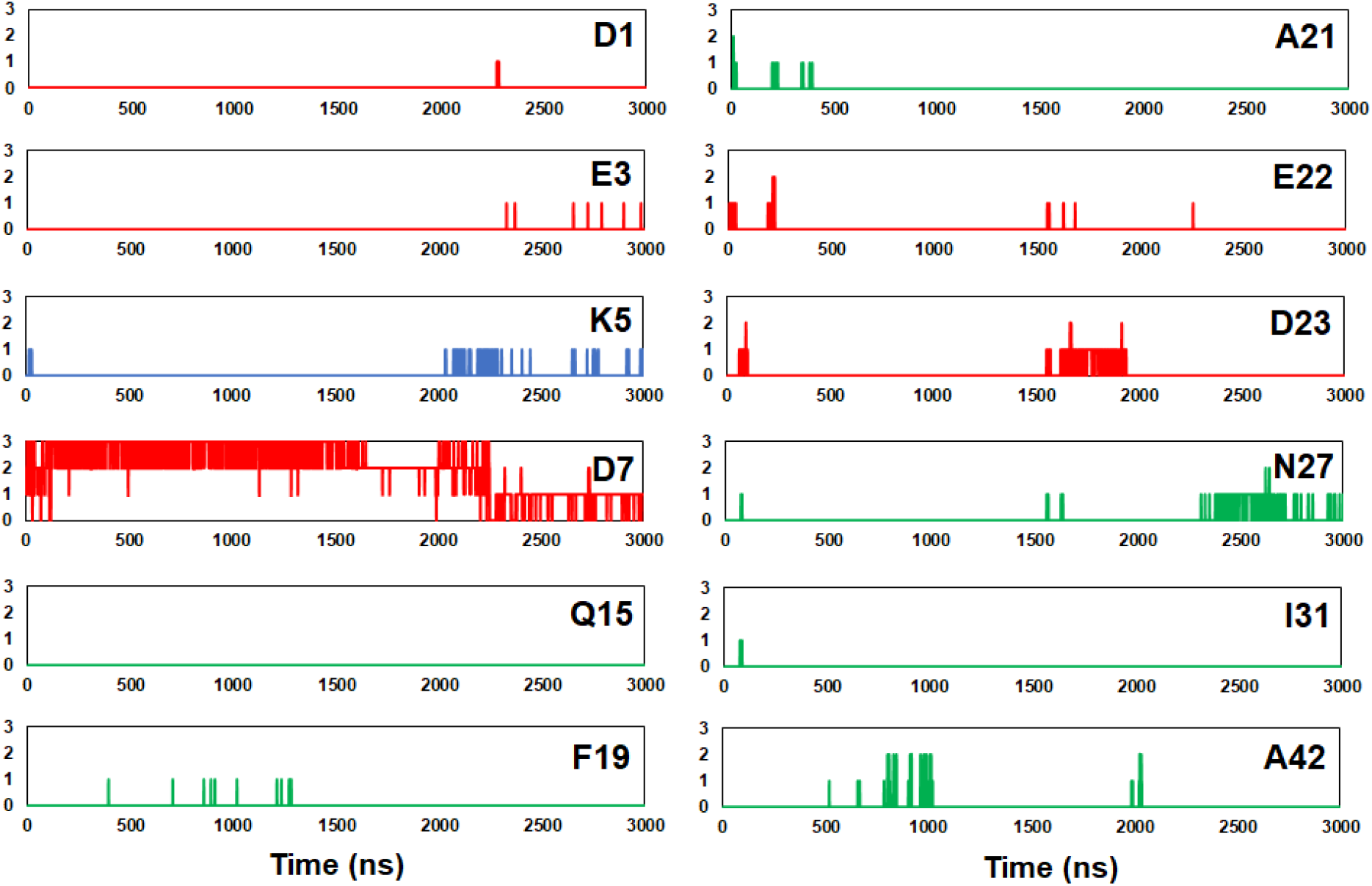
Hydrogen bonds between EGC and A*β*_42_ selected residues. Anionic, cationic, and neutral residues are shown in red, blue, and green, respectively.

**Figure S14.**
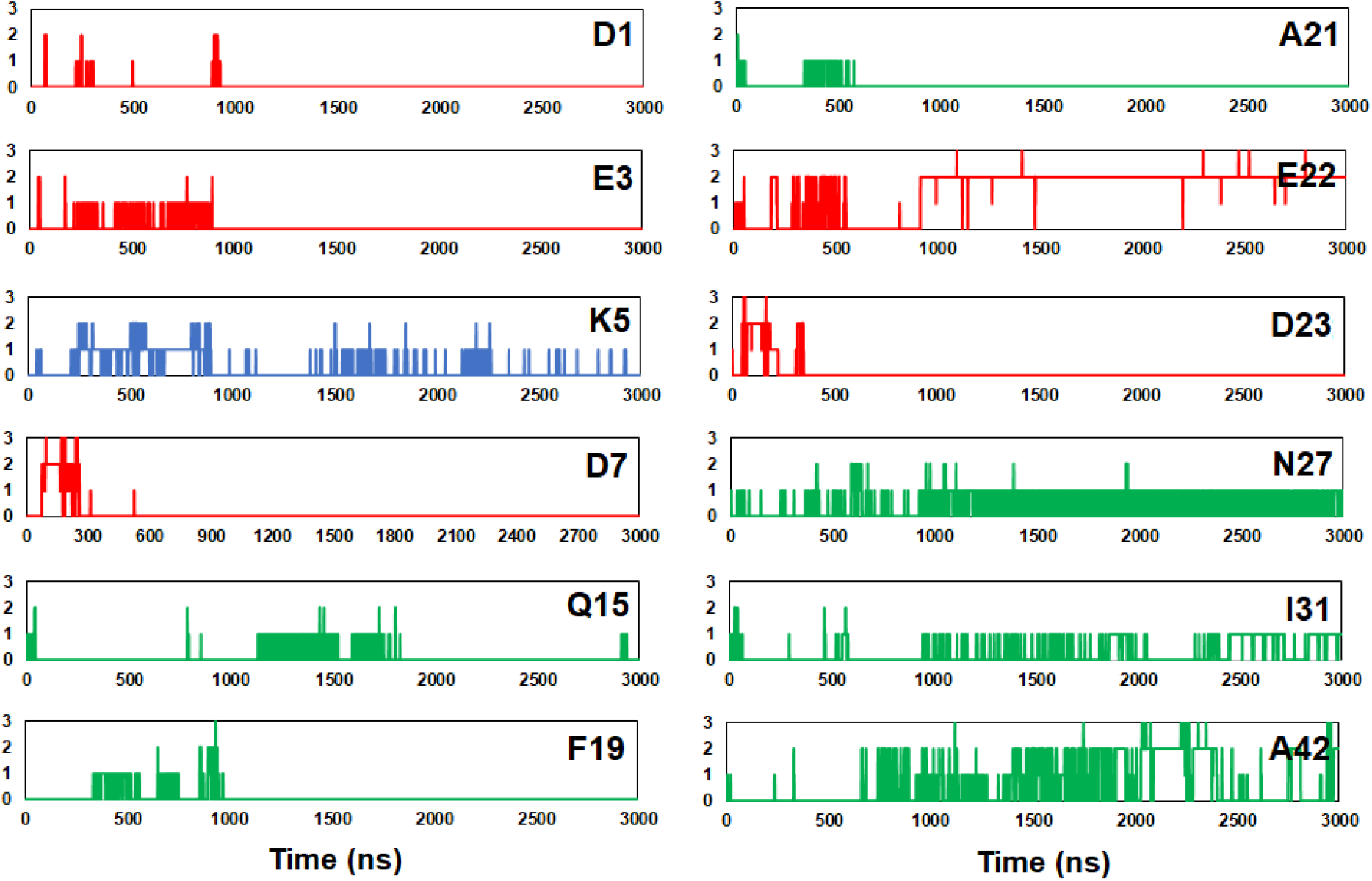
Hydrogen bonds between EGCG and A*β*_42_ selected residues. Anionic, cationic, and neutral residues are shown in red, blue, and green, respectively.

**Figure S15.**
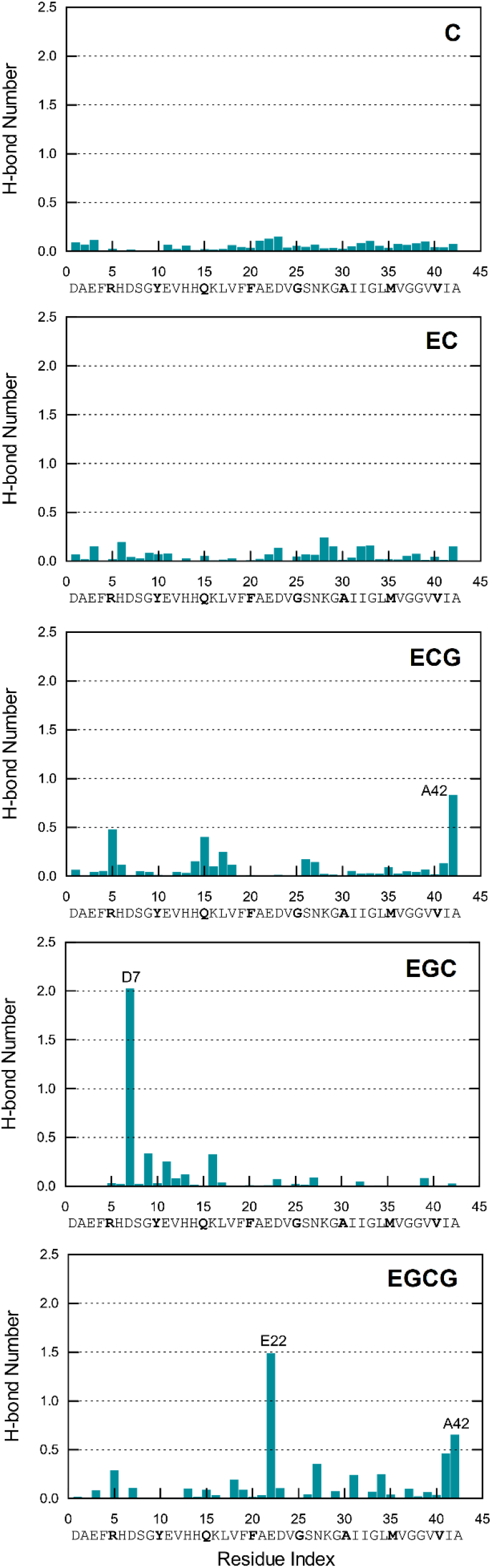
Average number of hydrogen bonds between A*β*_42_ amino acids and the ligands.

